# Reduction in Hepatic Phosphatidylcholine Biosynthesis Promotes MASH Through Copper Deficiency

**DOI:** 10.64898/2026.05.13.723926

**Authors:** Jaclyn E. Welles, James P. Garifallou, Michael V. Gonzalez, Dominic Santoleri, Feroza K. Choudhury, Gina M. DeNicola, Ryan W. Martin, Chang Jiang, Jaehee Kim, Gen Li, Yuichi Aki, Christopher J. Chang, David Li, Rebecca G. Wells, Yang Xiao, Jiayu Zhang, Mitchell A. Lazar, Donita C. Brady, Paul M. Titchenell

## Abstract

Metabolic dysfunction-associated steatohepatitis (MASH) is a progressive liver disease for which the mechanisms linking lipid dysregulation to fibrosis remain poorly defined. Hepatic phosphatidylcholine (PC) content is reduced in MASH, but how this alteration drives disease progression is unclear. Here, we identify a role for copper (Cu) homeostasis as a downstream effector of impaired PC biosynthesis. Using single-nucleus RNA sequencing in complementary genetic and dietary mouse models, we found that reduced hepatic PC is associated with marked depletion of hepatic Cu and a concomitant increase in circulating Cu, indicating disrupted Cu distribution. Mechanistically, PC depletion impaired plasma membrane localization of the high-affinity Cu transporter CTR1 (*SLC31A1*) in hepatocytes, limiting Cu uptake. In human hepatic stellate cells, Cu promoted fibrogenic activation, whereas suppression of Cu import or pharmacologic inhibition of MAPK signaling attenuated fibronectin deposition. *In vivo*, liver-directed Cu supplementation restored hepatic Cu levels and reduced steatosis but failed to improve fibrosis. In contrast, pharmacologic Cu chelation with bathocuproinedisulfonic acid (BCS) reduced fibrosis without affecting inflammation. Together, these findings identify Cu redistribution as a consequence of impaired PC biosynthesis and implicate Cu-dependent signaling in stellate cell activation, fibrogenesis and MASH pathogenesis.

**Graphical Abstract:** 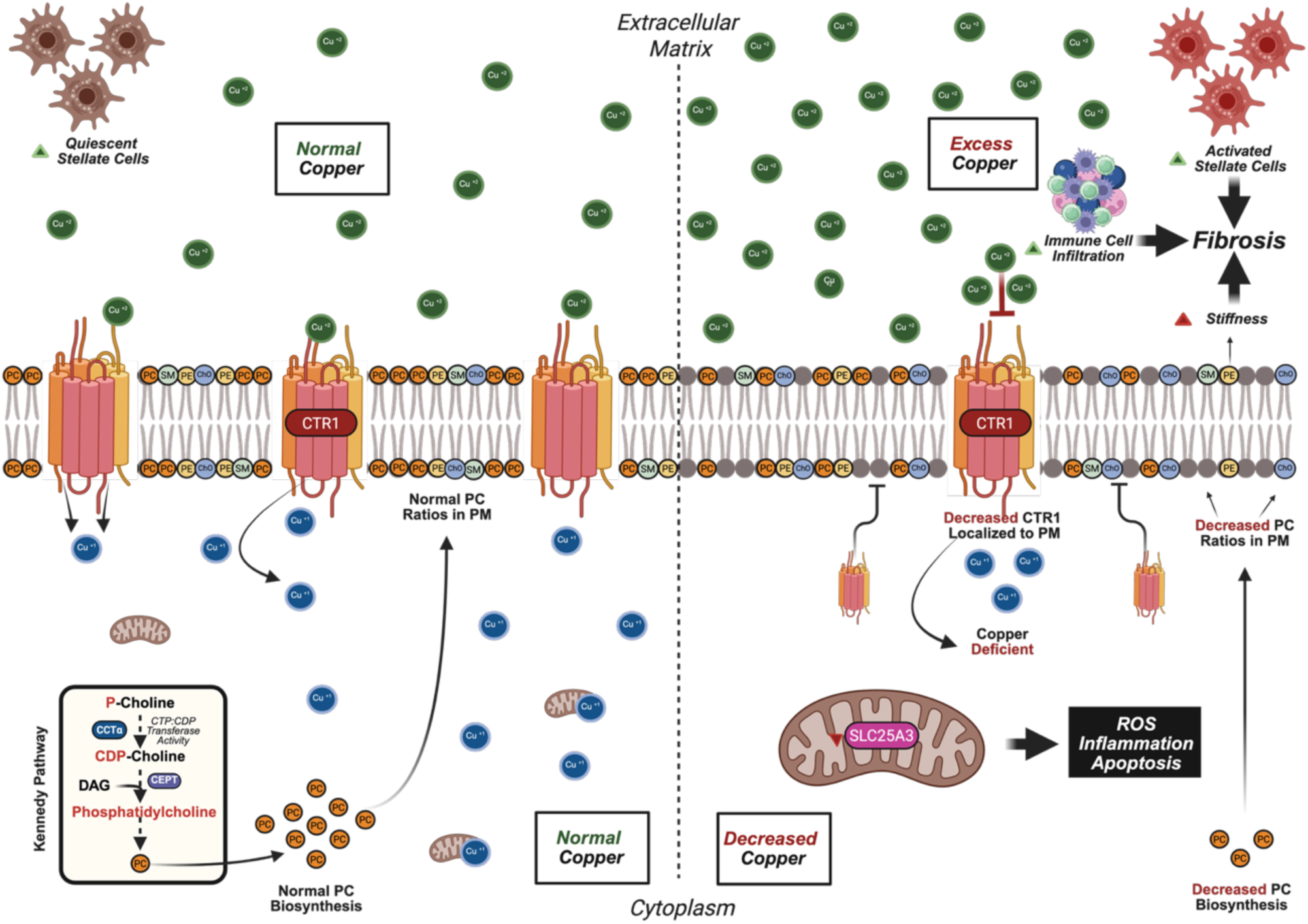

## Introduction

Metabolic dysfunction-associated steatohepatitis (MASH), formerly known as NASH, is a progressive liver disease arising from metabolic dysfunction-associated steatotic liver disease (MASLD) and is characterized by hepatocellular injury, inflammation, and fibrosis, representing a major cause of liver-related morbidity worldwide. Despite recent therapeutic advances, including the approval of resmetirom(1, 2), treatment options remain limited and have modest efficacy in improving fibrosis, and liver transplantation remains the only definitive therapy for advanced disease(3–5). These limitations underscore the need to define the mechanisms driving disease progression. Increased liver stiffness is a hallmark of MASH and correlates with disease severity and progression(3, 6–9). While extracellular matrix remodeling and inflammatory signaling molecules contribute to tissue stiffening (10–18), emerging evidence suggests that alterations in hepatic lipid composition also influence liver mechanics and fibrogenesis. In particular, changes in membrane lipid composition may impact cellular signaling pathways that drive stellate cell activation and fibrosis.

Alterations in hepatic lipid composition, including changes in cholesterol and phospholipids, are associated with increased inflammation and fibrosis across a spectrum of liver disease, including MASH(14, 19–29). Phosphatidylcholine (PC), one of the most abundant phospholipids in all tissues including the liver, plays a central role in membrane integrity and lipid metabolism(30, 31). Both experimental and human studies demonstrate that reduced hepatic PC biosynthesis promotes steatosis(22, 32–35), inflammation, and fibrosis(22, 32, 36–42). In mice, dietary and genetic disruption of PC synthesis induces features of MASLD and MASH, while in humans, variants in genes regulating PC biosynthetic pathways, including the Kennedy and PEMT pathways, are associated with disease susceptibility(38, 42–49). Consistent with these findings, multiple models of MASH, including Western diet-induced models(31, 50–53), the Gubra/AMLN-diet (GAN) (54–57) and carbon tetrachloride (CCL_4_) induced injury(50, 58, 59), exhibit progressive reductions in hepatic PC content, or PC ratios, as disease severity increases. Prior studies, including work from our group and others, have established a critical role for PC in very low-density lipoprotein (vLDL) secretion(31, 60–62), and neutral lipid accumulation (31, 32, 37, 63). Despite this, the exact mechanisms by which reduced PC biosynthesis drives fibrosis and inflammation remain poorly defined.

Here, we identify how impaired PC biosynthesis disrupts copper (Cu) homeostasis to contribute to MASH pathogenesis. Here, we identify disrupted copper (Cu) homeostasis as a consequence of impaired PC biosynthesis that contributes to MASH pathogenesis. Using complementary genetic and dietary models of reduced hepatic PC, we show that PC depletion impairs hepatocyte Cu uptake, leading to redistribution of Cu and altered cellular signaling. We further demonstrate that Cu promotes fibrogenic activation of hepatic stellate cells, whereas pharmacologic Cu chelation attenuates fibrosis *in vivo*. These findings define a mechanistic link between lipid remodeling and metal homeostasis in MASH and suggest that therapeutic strategies targeting site-specific Cu availability, restoring hepatocyte Cu to improve steatosis while limiting extracellular Cu to reduce fibrogenesis, may be required for effective treatment.

## Results

### Reduction in Hepatic PC Biosynthesis Drives MASH Pathology and Increases Liver Stiffness

To investigate the role of hepatic PC biosynthesis in MASH development, we employed complementary genetic and dietary models of PC deficiency. For the genetic model, *PCYT1A* (CCT-α) floxed mice consumed a chow diet and we administered AAV8-TBG-CRE to induce hepatocyte-specific knockout of *PCYT1A* (L-CCTα-KO). For the dietary model, mice consumed a low-methionine, choline-deficient high-fat diet (LMCD-HFD) for 4 weeks to impair *de novo* PC synthesis via both the PEMT and Kennedy pathways (31, 62).

Consistent with prior studies, both models exhibited hepatic micro- and macro-steatosis, collagen deposition, and fibrosis (Figure 1A) (58, 59, 63). Despite reductions in overall body weight (Figure 1B), Both experimental groups showed significantly increased liver weight compared with controls (Figure 1C). They also showed elevated markers of liver injury, including aspartate aminotransferase (AST) and alanine aminotransferase (ALT) (Figure 1, D and E). In parallel, expression of pro-fibrogenic genes (*Saa1, Col1a1, Timp1*) and pro-inflammatory markers, including C-reactive protein (CRP) and inflammatory cytokines (*Ccl3, Ptgs2, Cxcl1*), increased (Figure 1F and Supplemental Figures 1A and B). Lipogenic genes *Fads1* and *Fads3* also significantly increased in mouse MASH livers compared to control (Supplemental Figure 1C).

**Figure 1.**
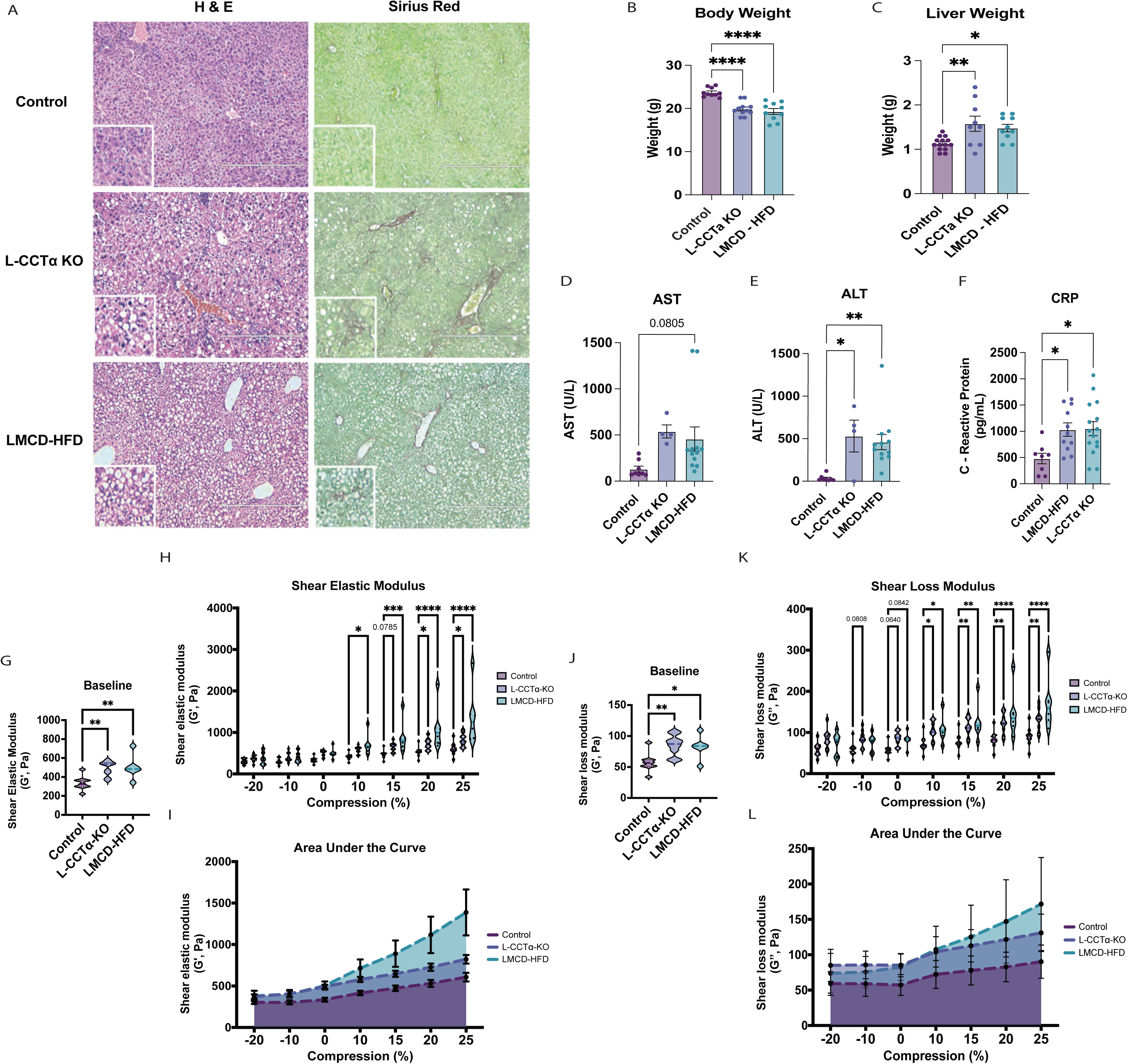
Impaired Hepatic PC Biosynthesis Promotes MASH Pathology and Increases Liver Stiffness. **(A)** Representative histology showing H&E and fibrosis via Sirius red staining in control vs. genetic (L-CCTα-KO), and dietary (LMCD-HFD) mouse MASH livers. **(B)** Total body weight (BW) in mouse MASH livers compared to control. n = 10-12. **(C)** – Liver weight (LW) in mouse MASH livers compared to control. n = 10-12. **(D)** - AST, **(E)** - ALT, and **(F)** - C-Reactive protein (CRP) in mouse MASH models compared to control. n = 8-15. **(G) – (L)** – Stiffness and viscoelasticity across physiological ranges of compression in mouse MASH livers compared to control. n = 6-8. Data are shown as mean ± SEM. *P < 0.05; **P < 0.01; ***P < 0.001. Scale bars: 400 μm.

Because increased liver stiffness is a hallmark of MASH progression (3, 66–72), we next assessed the mechanical properties of livers from both models. Rheological measurements revealed a significant increase in stiffness and altered viscoelasticity, as indicated by elevated shear elastic (Figure 1, G and H) and shear loss (Figure 1, I-J) moduli. Together, these findings demonstrate that impaired hepatic PC biosynthesis is sufficient to drive key features of MASH, including increased liver stiffness, suggesting that PC-dependent lipid remodeling contributes directly to the mechanical changes associated with MASH progression.

### PC Deficiency Alters Hepatic Cell States and Membrane-Associated Pathways in MASH

To further define the cellular and molecular consequences of impaired phosphatidylcholine (PC) biosynthesis, we performed single-nucleus RNA sequencing (snRNA-seq) on livers from chow-fed control, hepatocyte-specific *PCYT1A* knockout (L-CCTα-KO), and LMCD-HFD-fed mice. Unsupervised clustering identified major hepatic cell populations, including hepatocytes (HEP), hepatic stellate cells (HSCs), liver sinusoidal endothelial cells (LSECs), Kupffer cells (KCs), and cholangiocytes (CHO) (Figure 2, A-C and Supplemental Figure 2A). In addition, we identified a distinct cluster of nuclei characterized by elevated Diaph3 (mDia3) expression (UNK). Cell type annotations were confirmed using previously published mouse and human liver single-cell datasets (Supplemental Figure 2B and C)(73).

**Figure 2.**
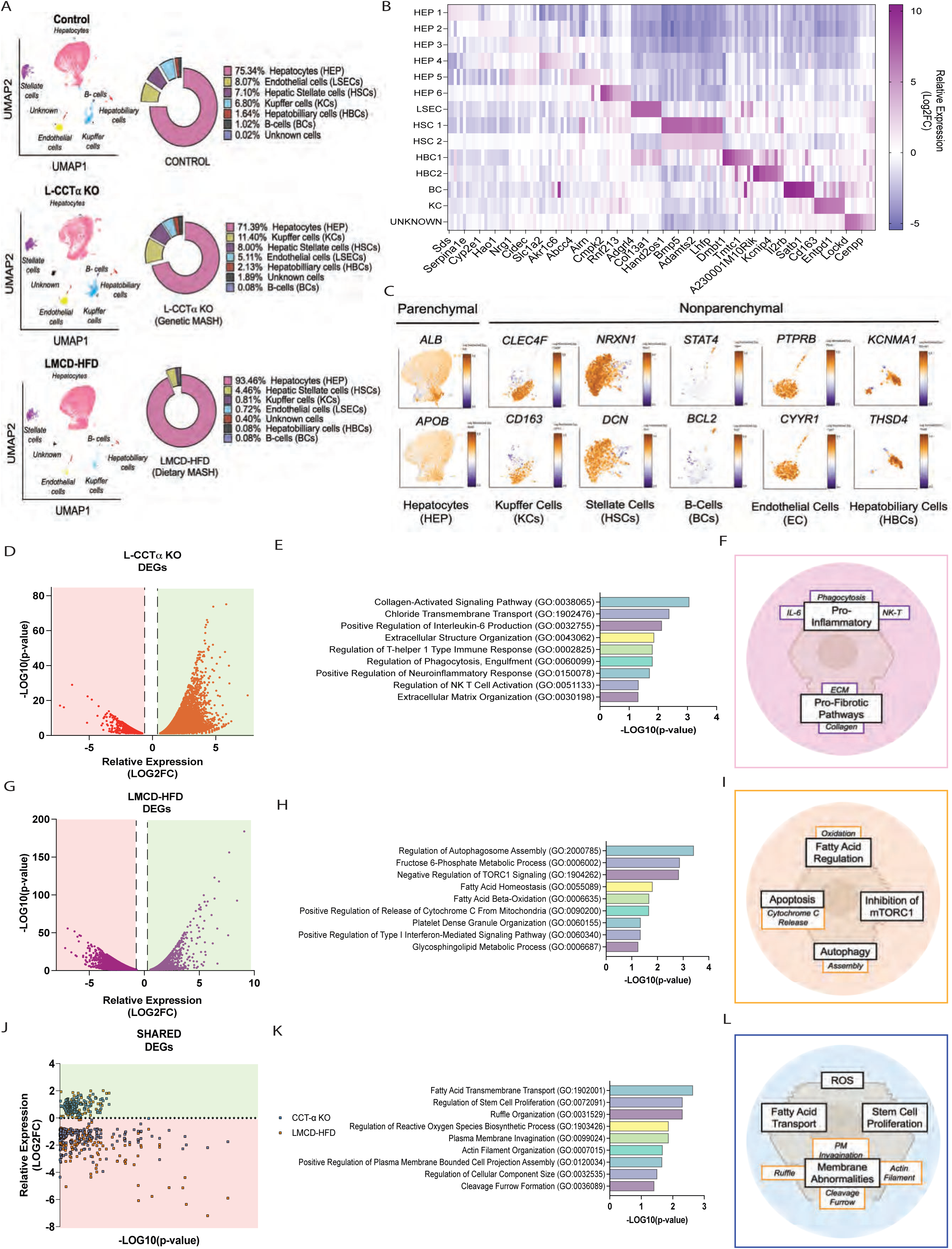
Single-Nucleus RNA-Seq Reveals PC-Dependent Changes in Hepatic Cell States and Membrane Pathways in MASH. **(A)** – SEURAT Cluster analysis between cells enriched in mouse MASH livers compared to control. **(B)** – Heat map of consolidated DEG identified between clusters in whole livers. **(C)** – Annotation of specific clusters identified in Control and MASH livers. **(D)** – Volcano plot of DEGs identified in genetic MASH livers (total) compared to control livers. **(E)** – KEGG Pathway analysis of DEGs enriched in genetic MASH livers. **(F)** – Schematic of KEGG Profile in genetic MASH. **(G)** – Volcano plot of DEGs identified in dietary MASH livers (total) compared to control livers. **H)** - KEGG Pathway analysis of DEGs enriched in dietary MASH livers. **(I)** - Schematic of KEGG Profile in dietary MASH. **(J)** – Volcano plot of DEGs shared in both MOUSE MASH livers (total) compared to control. **(K)** – KEGG Pathway analysis of shared DEGs enriched in both the genetic and dietary MASH livers. **(L)** - Schematic of KEGG Profile from both MASH models.

Pathway analysis of differentially expressed genes revealed enrichment of pro-fibrogenic pathways in the genetic MASH model (Figure 2, D-F), and lipid metabolic pathways in the dietary model (Figure 2, G-I). Notably, genes shared between both models were significantly enriched for membrane-associated processes, including cleavage furrow formation, plasma membrane invagination, and ruffle organization (Figure 2, J-L), suggesting widespread alterations in membrane dynamics.

Given the importance of hepatic zonation in liver function and its disruption in advanced fibrosis (65, 72–75). We next assessed zonation markers. Both MASH models displayed reduced expression of the periportal marker *Sds*, whereas only the genetic model selectively decreased the pericentral marker *Cyp2e1* (Supplemental Figure 2, D-F). These findings indicate that impaired PC biosynthesis disrupts hepatocyte membrane organization and contributes to loss of hepatic zonation, suggesting that PC-dependent membrane remodeling is a key determinant of hepatocyte organization and function in MASH.

### PC Deficiency Impairs Hepatocyte Cu Uptake and Disrupts Cu Homeostasis

To determine how impaired PC biosynthesis alters hepatocyte function, we analyzed snRNA-seq data from hepatocyte populations across control, genetic (L-CCTα-KO), and dietary MASH models. Of the 26,803 hepatocyte nuclei identified, 9,854 were derived from control livers, 7,017 from genetic MASH livers, and 9,932 from dietary MASH livers (Figure 3A). Approximately 10,239 genes were differentially expressed between genetic and dietary MASH hepatocytes when compared to control (Supplemental Figure 3A). Differential gene expression analysis revealed significant enrichment of pathways associated with metal ion handling, including response to metal ions and transition metal ion binding, in hepatocytes from both mouse MASH models as well as human single-cell RNA sequencing (scRNA-seq) MASH datasets (Figure 3, B-E and Supplemental Figure 3C and D).

**Figure 3.**
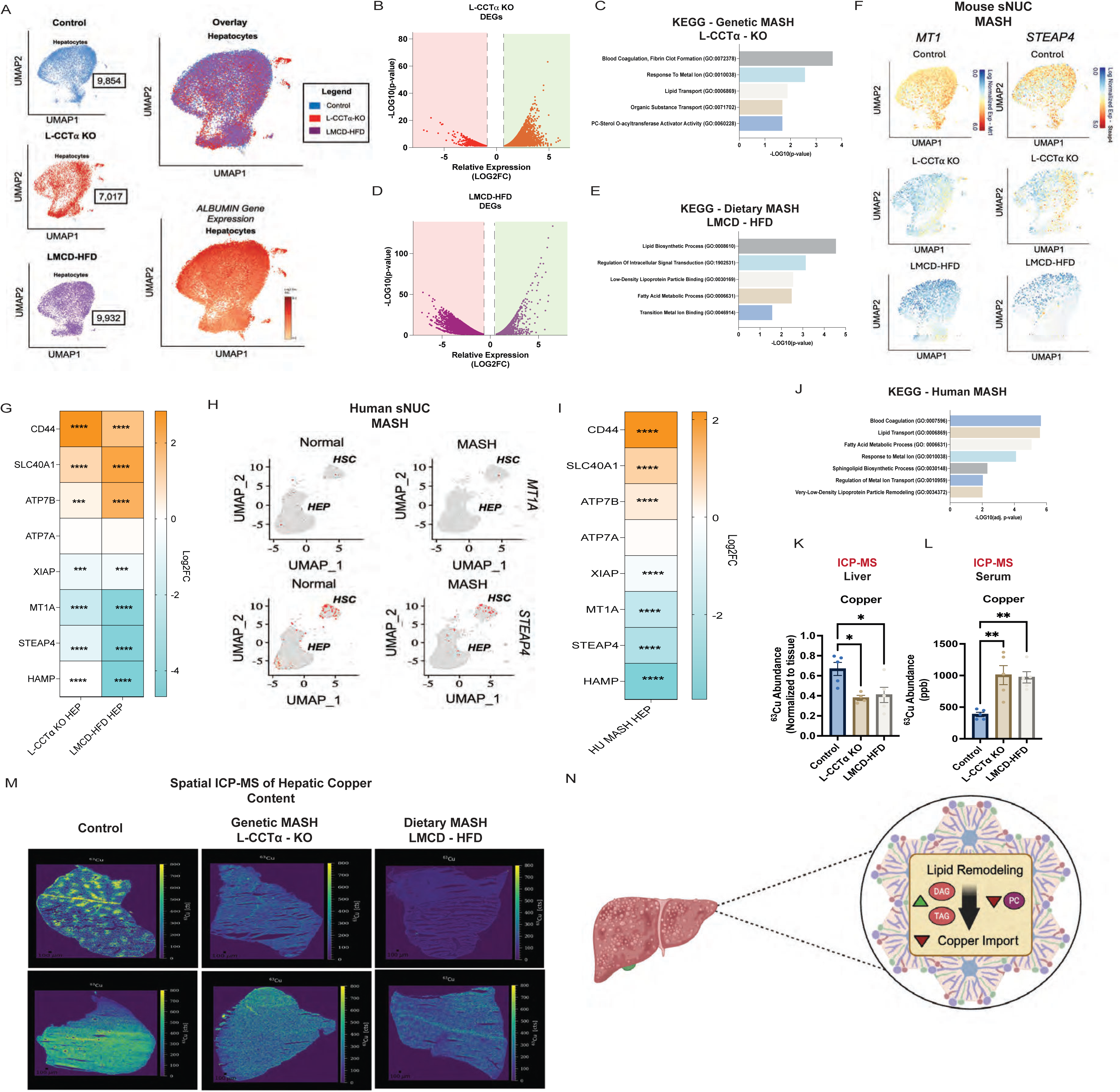
PC Deficiency Disrupts Hepatic Copper Homeostasis and Reduces Hepatocyte Copper Content in MASH. **(A)** – U-Maps illustrating hepatocytes from control vs mouse MASH livers. Albumin gene expression was used as a “positive control”. **(B)** – Volcano plot showing DEGs between control and genetic mouse MASH hepatocyte clusters. **(C)** – KEGG pathway analysis of DEGs between control and genetic mouse MASH hepatocyte clusters. **(D)** - Volcano plot showing DEGs between control and dietary mouse MASH hepatocyte clusters. **(E)** – KEGG pathway analysis of DEGs between control and dietary mouse MASH hepatocyte clusters. **(F)** – U-Maps showing gene expression overlay of Cu-specific genes in control vs. mouse MASH hepatocytes. **(G)** – Heat map of Metal regulated genes differentially expressed between control and mouse MASH hepatocytes. **(H)** – U-Maps showing gene expression overlay of Cu-specific genes in control and human MASH hepatocytes. **(I)** - Heat map of Metal regulated genes differentially expressed between control and human MASH hepatocytes. **(J)** – KEGG pathway analysis of DEGs between control and human MASH hepatocyte clusters. **(K)** – ICP-MS showing normalized ^63^Cu in liver tissue from control and mouse MASH livers. n = 5-6. **(L)** - ICP-MS showing normalized ^63^Cu in the serum of control and mouse MASH livers. n = 5-6. **(M)** – Spatial ICP-MS showing reduction in total copper accumulated in mouse MASH livers compared to control. **(N)** – Schematic of summarized data. Data are shown as mean ± SEM. *P < 0.05; **P < 0.01; ***P < 0.001; NS, no significance, P > 0.05

Consistent with these pathway-level changes, expression of Cu-associated genes, including *MT1*, *SOD1*, and *STEAP4*, was reduced in hepatocytes from both mouse and human MASH samples (Figure 3, F-J). Notably, these changes occurred without significant alterations in expression of the high-affinity Cu importer *SLC31A1* (CTR1). In contrast, expression of the hepatocyte Cu exporter ATP7B was increased, suggesting a shift in cellular Cu handling (Supplemental Figure 3B).

To determine whether these transcriptional changes were associated with altered Cu levels, we quantified hepatic and circulating Cu by inductively coupled plasma mass spectrometry (ICP-MS). Both genetic and dietary MASH models exhibited a significant reduction in hepatic Cu content (∼40%) compared with controls (Figure 3, K and L), a finding further supported by spatial ICP-MS imaging (Figure 3M). Together, these data demonstrate that impaired PC biosynthesis disrupts hepatocyte Cu homeostasis, consistent with impaired Cu uptake and altered Cu distribution (Figure 3N).

### PC Deficiency Promotes Pro-fibrogenic Activation of Hepatic Stellate Cells in MASH

Given the central role of HSCs in fibrogenesis, we next assessed whether hepatocyte-specific reduction in PC is sufficient to alter HSC transcriptional states. snRNA-seq revealed expansion of HSC populations in both genetic (n = 789) and dietary (n = 420) MASH livers compared with controls (Figure 4A). These HSC clusters exhibited increased expression of pro-fibrogenic genes, including *Pth1r*, *Tgfbi*, *Col1α1*, and *Loxl2* (Figure 4, B and C), consistent with activation. In contrast, quiescent HSC populations characterized by expression of *Itgb3* and *Tnfrsf11b* were observed predominantly in control livers and not MASH models, a finding further supported by decreased expression of the quiescence-associated marker CYGB (Supplemental Figure 4, A and B).

**Figure 4.**
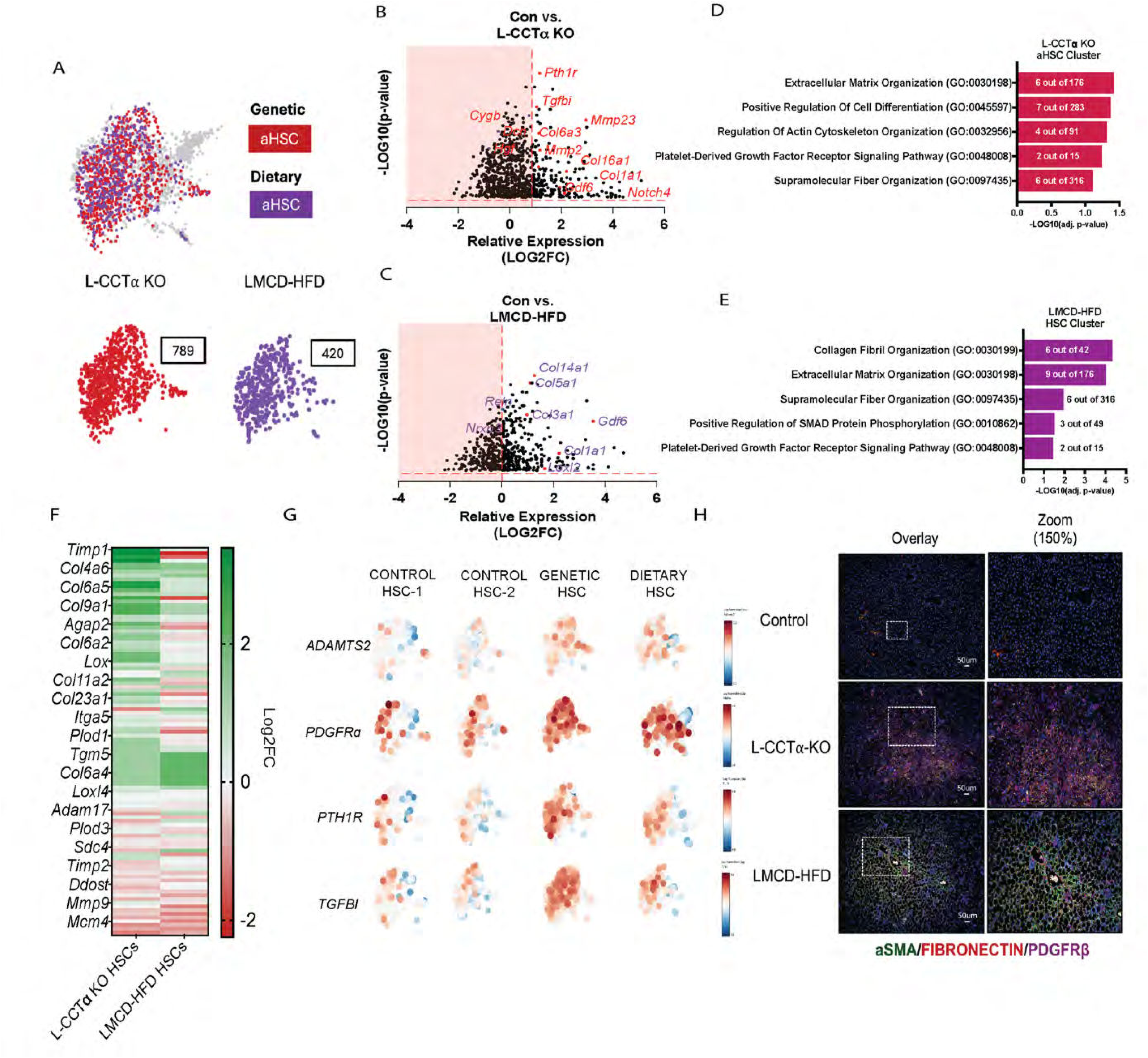
PC Deficiency Promotes Pro-fibrogenic Activation of Hepatic Stellate Cells in MASH. **(A)** – Louvain cluster analysis showing hepatic stellate cells (HSCs) identified in each mouse MASH model. **(B-C)** – Volcano plots of differentially expressed genes (DEGs) in HSCs from genetic vs. dietary mouse MASH model. **(D)** – KEGG Pathway analysis of DEGs in HSC from genetic mouse MASH livers compared to total HSCs enriched in dataset. **(E)** - KEGG Pathway analysis of DEGs in HSC from dietary mouse MASH livers compared to total HSCs enriched in dataset. **(F)** – Heat map of DEGs associated with collagen crosslinking and fibrosis in HSCs from mouse MASH livers compared to control. **(G)** – U-map showing gene expression overlay of DEGs associated with collagen crosslinking and fibrosis in HSCs from mouse MASH livers compared to control. **(H)** – Immunofluorescence (IF) stains of HSCs in control vs. mouse MASH livers.

Pathway analysis of the 471 differentially expressed genes shared between genetic and dietary MASH HSCs demonstrated enrichment of pathways associated with extracellular matrix (ECM) organization, PDGFRβ signaling, supramolecular fiber organization, and SMAD phosphorylation (Figure 4, D and E), consistent with fibrogenic activation. Genes involved in collagen crosslinking and extracellular matrix deposition were also significantly upregulated (Figure 4F and G). Consistent with these transcriptional changes, immunofluorescence analysis revealed expansion of αSMA- and fibronectin (FN1)-positive cells in MASH livers compared with controls, indicating increased HSC activation and matrix deposition (Figure 4H). Notably, similar patterns of αSMA and FN1 localization were observed in fibrotic regions of human liver samples (Supplemental Figure 4, C and D), supporting the translational relevance of these findings and consistent with a role for impaired PC biosynthesis in promoting stellate cell activation and extracellular matrix remodeling in MASH.

### PC Depletion Disrupts CTR1 Localization and Impairs Hepatocyte Cu Uptake

To determine whether reduced hepatic Cu content reflects impaired Cu import, we assessed CTR1 localization in HUH-7 cells following knockdown of *PCYT1A*. Immunofluorescence analysis using a pre-conjugated CTR1 probe revealed reduced plasma membrane localization of CTR1 in CCTα-deficient cells under low-Cu conditions, including BCS treatment or serum-free media (Figure 5A). In parallel, altered localization and expression of the Cu exporter ATP7B were observed following *PCYT1A* knockdown in CuSO₄-treated cells (Figure 5B).

**Figure 5.**
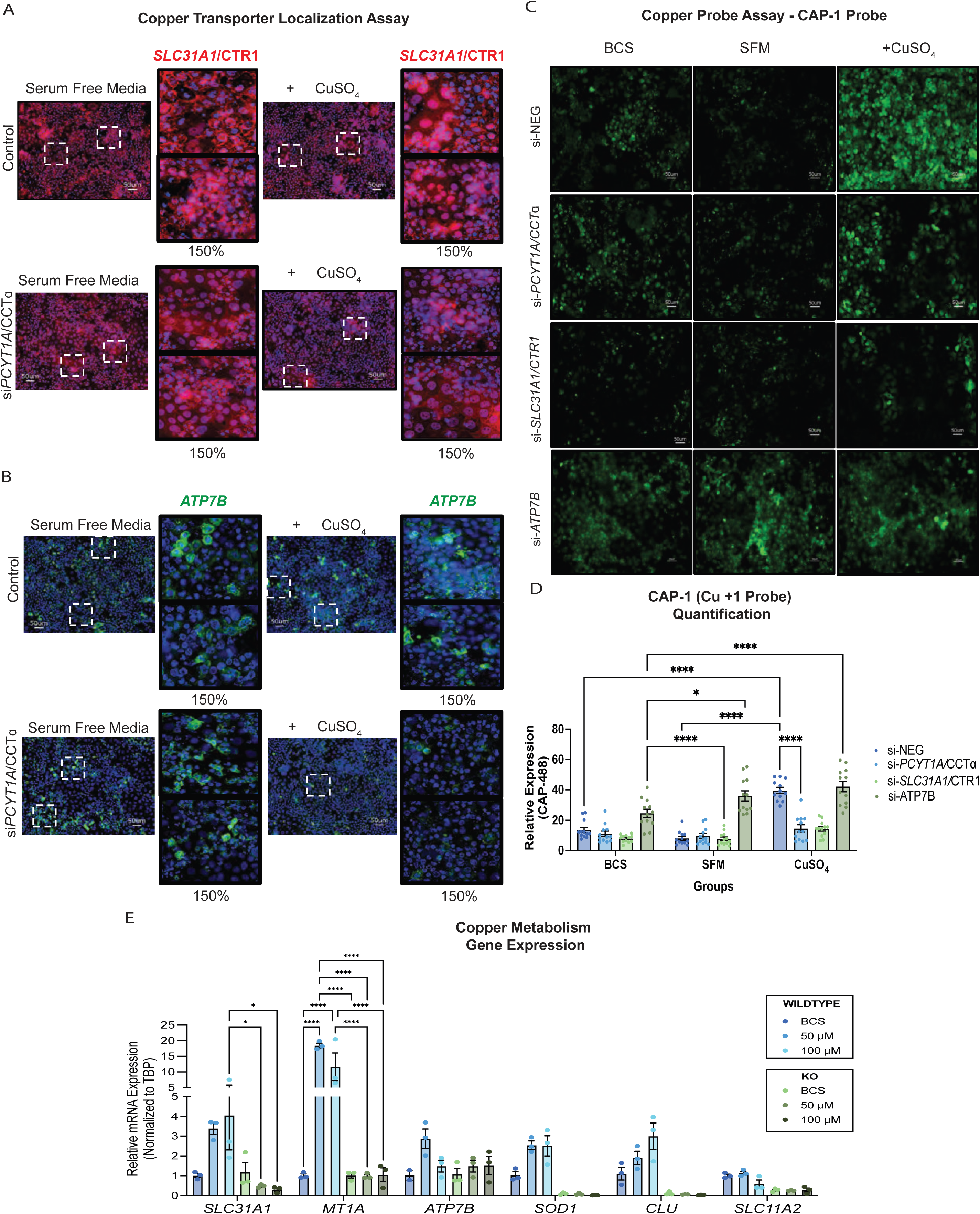
PC Depletion Disrupts CTR1 Localization and Impairs Hepatocyte Copper Uptake. **(A)-(B)** – Immunofluorescence (IF) showing CTR1 **(A)** and ATP7B **(B)** localization in siRNA-negative control, siRNA-PCTY1A treated HUH-7 cells with or without CuSO4. n = 6-8. **(C)** – IF showing Cu import using CAP-1, a Cu^+1^ specific probe that emits a fluorescence at 488 in the presence of copper in HUH-7 cells transfected with siRNA against *PCYT1A*, *SLC31A1*/CTR1 (importer) or *ATP7B* (exporter). n = 6-8. **(D)** – Quantification of CAP-1 Cu-Probe + cells/signal in HUH-7 cells +/- *PCYT1A*, CTR1, or ATP7B after being treated with either Serum Free Media (SFM), 500 μM BCS, or 50 μM CuSO_4_ for 6-hr. n = 12. **(E)** RT-qPCR of HUH-7 cells in the presence or absence of either siRNA-Negative Control or siRNA-*PCTY1A*, after being treated with either Serum Free Media (SFM), 500 μM BCS, 50 μM, or 100 μM CuSO_4_for 6-hr. n = 3-6 per group. Scale bars: 50 μm. Data are shown as mean ± SEM. *P < 0.05; **P < 0.01; ***P < 0.001.

To directly assess Cu uptake, we measured intracellular Cu⁺ using the fluorescent probe CAP-1(78). Knockdown of *PCYT1A* significantly reduced CAP-1 fluorescence compared with control cells, indicating impaired Cu import (Figure 5, C and D). As expected, silencing *SLC31A1* reduced Cu⁺ accumulation, whereas knockdown of *ATP7B* increased intracellular Cu⁺ levels, validating the assay. Consistent with these findings, induction of Cu-responsive genes, including *MT1A*, *MT1B*, and *SOD1*, was blunted in CCTα-deficient cells following CuSO₄ treatment (Figure 5E).

To determine whether altered membrane lipid dynamics drove these effects, we disrupted phosphatidylcholine incorporation into the plasma membrane by silencing phosphatidylcholine transfer protein (*PC-TP*) and observed similar defects in CTR1 localization and ATP7B response, supporting a role for membrane PC in regulating Cu transport (Supplemental Figure 5, A-D).

Western blot analysis demonstrated a dose-dependent reduction in CTR1 protein levels following Cu treatment, which was not altered by *PCYT1A* knockdown. In contrast, ATP7B protein exhibited altered electrophoretic mobility in CCTα-deficient cells, suggesting potential changes in protein processing or conformation. In addition, Cu-induced ERK phosphorylation was reduced in CCTα-deficient cells, further supporting impaired Cu-dependent signaling (Supplemental Figure 5, E-F). Dose-dependent increase in Cu-binding protein, metallothionein 1A and 1B (*MT1A*/*MT1B*), was also observed in HUH-7 cells (Supplemental Figure 5, G). Together, these data demonstrate that PC-dependent membrane remodeling is required for proper CTR1 localization and Cu import, providing a mechanistic basis for hepatocyte Cu deficiency in MASH.

### Cu Promotes Fibrogenic Activation of Hepatic Stellate Cells in a MAPK-Dependent Manner

Given the impairment in hepatocyte Cu uptake, we next asked whether altered Cu availability promotes fibrogenic activation of nonparenchymal cells, particularly HSCs. To test this, we treated human LX-2 stellate cells with CuSO₄ and either a low- or high-dose of TGFβ. Cu treatment alone modestly increased expression of pro-fibrogenic genes, including *COL1A1*, *COL6A3*, *TIMP1*, *PDGFRβ*, and *FN1*. However, Cu markedly enhanced TGFβ-induced transcriptional responses, indicating an additive effect on fibrogenic activation (Figure 6A).

**Figure 6.**
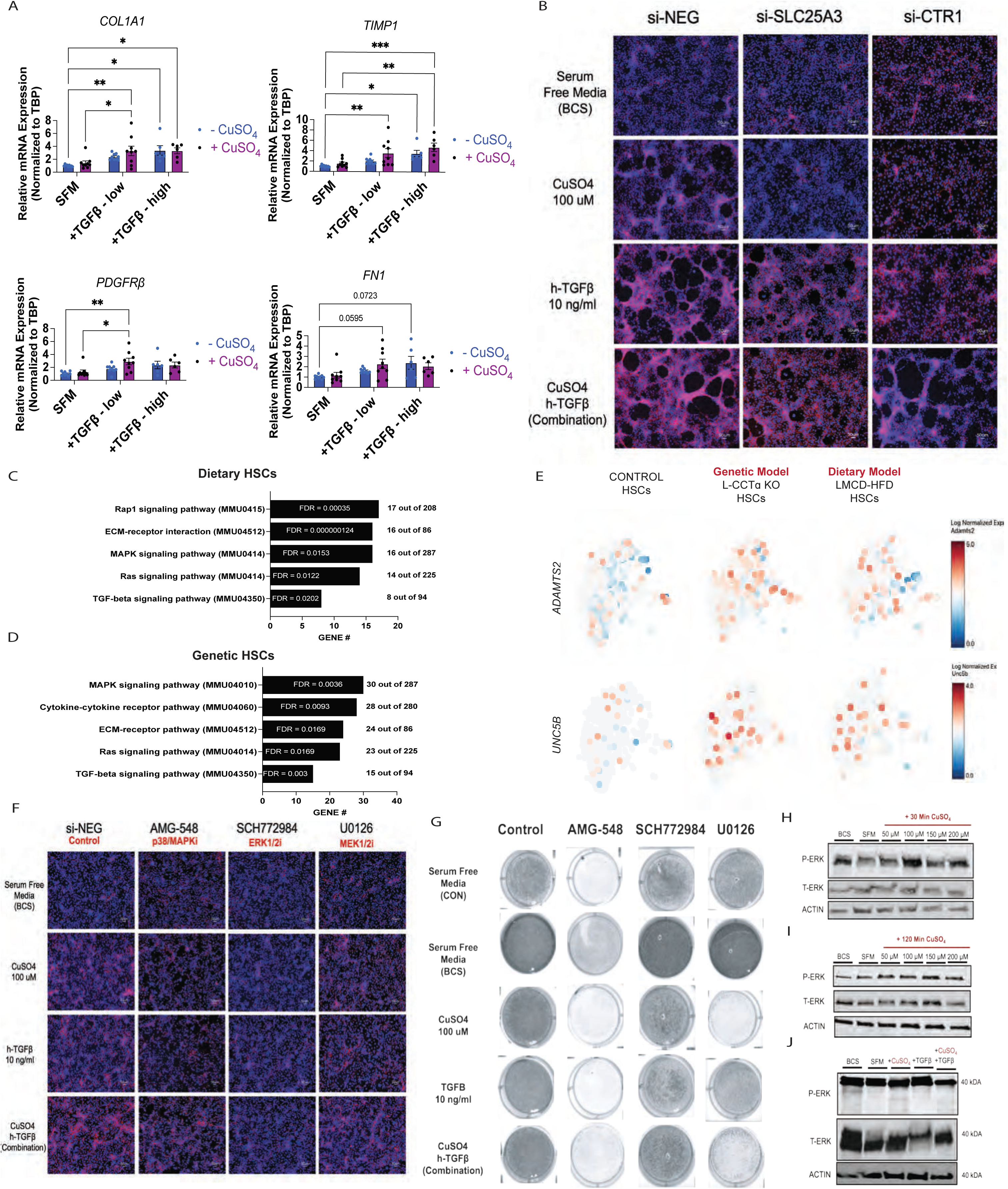
Copper Promotes Fibrogenic Activation of Hepatic Stellate Cells in a MAPK-Dependent Manner. **(A)** – RT-qPCR of pro-fibrogenic gene expression in response to 24-hr 100 μM CuSO_4_ +/- low TGFβ (10 ng/mL), or high TGFβ (100 ng/mL) in LX-2 cells. **(B)** – IF showing fibronectin deposition following transfection of either siRNA-*SLC31A1*/CTR1 (PM), or siRNA-*SLC25A3* (Mito). **(C)** – STRING Pathway analysis of stellate cell cluster in genetic model. **(D)** – STRING Pathway analysis of stellate cell cluster in dietary model. **(E)** – Louvain cluster heat map of genes upregulated in stellate cell cluster, associated with MAPK pathway. **(F)** – IF showing fibronectin deposition in LX-2 cells pre-treated with various MAPK inhibitors for 6-hr. **(G)** - Crystal violet stain of LX-2 treated with various inhibitors with or without TGFβ or 100 μM CuSO_4_ for 24-hrs. **(H-J)** – Western blots showing dose dependent changes in ERK1/2 phosphorylation in LX-2 cells treated with either just 100 μM CuSO_4_ or 100 μM CuSO_4_ and TGFβ (10 ng/mL). Data are shown as mean ± SEM. *P < 0.05; **P < 0.01; ***P < 0.001. Scale bars: 50 μm.

Consistent with these findings, Cu treatment increased fibronectin deposition and extracellular matrix formation, as measured using an FN1-conjugated fluorescent probe, effects that were further amplified by TGFβ (Supplemental Figure 6, A and B). To determine whether intracellular Cu availability is required for fibrogenesis, we disrupted distinct pathways of Cu import in LX-2 cells. Silencing the plasma membrane Cu transporter *CTR1* (*SLC31A1*) reduced fibronectin deposition, consistent with impaired Cu uptake. Reductions in pro-fibrogenic gene expression was also observed in response to Cu and TGFβ (Supplemental Figure 6C). Notably, knockdown of the mitochondrial Cu transporter *SLC25A3* similarly attenuated fibronectin deposition (Figure 6B), indicating that intracellular Cu delivery to at least the mitochondrial compartment is required for fibrogenic activation. However, contrary to CTR1 knockdown, *SLC25A3* knockdown did not prevent Cu and TGFβ-induced pro-fibrogenic gene expression (Supplemental Figure 6D). This data suggest that copper may affect fibrogenesis through a mechanism independent of transcription such as, mitochondria-induced oxidative stress, cell death (cuproptosis), or the production of reactive oxygen species (ROS).

Pathway analysis of HSC clusters from both genetic and dietary MASH models revealed enrichment of MAPK signaling pathways (Figure 6, C and D), with upregulation of MAPK-associated genes such as *Adamts2* and *Unc5b* (Figure 6E). Pharmacologic inhibition of MAPK signaling using p38 (AMG-548), MEK1/2 (U0126), or ERK1/2 (SCH772984) inhibitors significantly reduced fibronectin deposition and suppressed pro-fibrogenic gene expression in response to Cu and TGFβ (Figure 6F and Supplemental Figure 6E). Cell viability stains also showed Cu-specific lethality in cells treated with U0126, suggesting that in stellate cells, MEK1/2 activity is required in the presence of Cu (Figure 6G). Consistent with this, Cu treatment increased ERK1/2 phosphorylation in LX-2 cells (Figure 6, H-J). Together, these findings demonstrate that Cu promotes fibrogenic activation of HSCs through MAPK-dependent signaling pathways.

### Hepatic Cu Supplementation Reduces Steatosis but Not Fibrosis in MASH

To assess the therapeutic potential of restoring hepatic Cu levels, we administered the liver-directed Cu supplement [CuS; Gal-Cu (GTSM)] (79), to mice subjected to a 4-week LMCD-HFD challenge (Figure 7A). ICP-MS analysis confirmed that CuS restored hepatic Cu content in MASH livers to levels comparable to controls (Figure 7, B and C). Restoration of hepatic Cu was associated with significant improvements in lipid metabolism, including reduced liver weight and decreased hepatic triglyceride accumulation (Figure 7, D-F). As expected, CuS - treatment did not rescue any other serum or hepatic metals dysregulated in MASH livers (Supplemental Figure 8). Histological analysis further demonstrated reduced steatosis in CuS-treated mice, accompanied by a shift toward lower steatosis grades, including a marked reduction in stage 3 steatosis and enrichment of intermediate stages (Figure 7, G and H). In contrast, CuS treatment did not improve fibrosis. Histological scoring revealed no significant reduction in fibrosis severity (Figure 7I), and expression of pro-fibrogenic and pro-inflammatory genes remained elevated (Figure 7, J and K). No changes in Cu transport genes were observed (Figure 7L). Together, these data demonstrate that restoration of hepatocyte Cu is sufficient to improve steatosis but is insufficient to resolve fibrosis or inflammation in MASH. These findings suggest that hepatic Cu repletion alone does not fully correct the pathogenic effects of altered Cu homeostasis and support a model in which compartment-specific Cu availability differentially regulates steatosis and fibrogenesis (Figure 7M).

**Figure 7.**
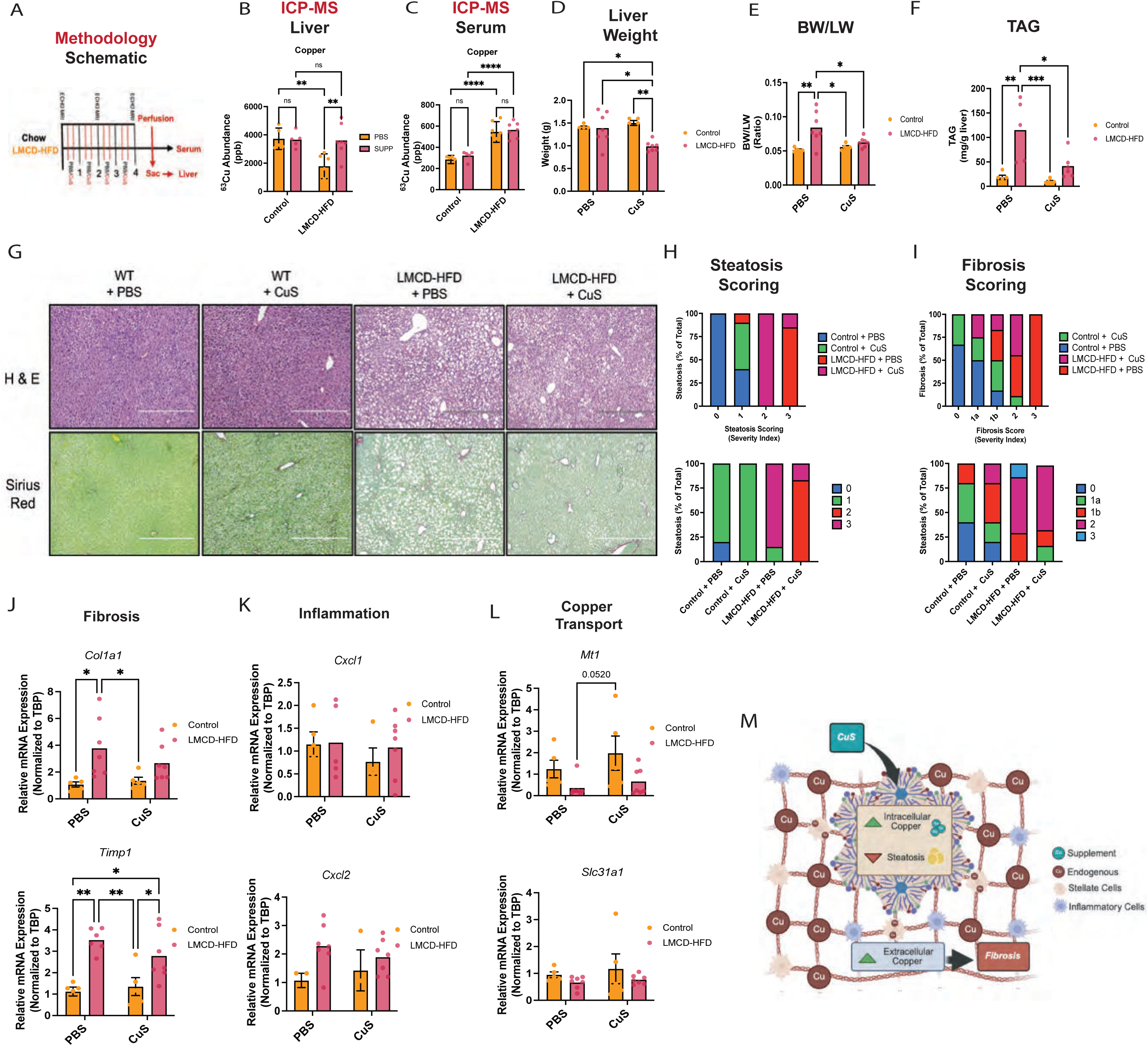
Hepatic Copper Supplementation Reduces Steatosis but Not Fibrosis in MASH. **(A)** – Schematic of Study Design and Model. ICP-MS showing normalized ^63^Cu in the liver **(B)** or serum **(C)** of control vs. dietary mouse MASH mice post PBS or CuS administration for 4-wks. Liver weight (D), and liver weight normalized to toal body weight (BW) **(E)** in control vs. dietary mouse MASH mice post PBS or CuS administration n = 5-7. **(F)** – Triglyceride (TAG) assay quantifying liver fat accumulation in livers post PBS or CuS administration n = 5-6. **(G)** – Representative Histology. **(H)** – Blind steatosis, and **(I)** fibrosis scoring, in control vs. dietary mouse MASH mice post PBS or CuS administration. **(J-L)** – RT-qPCR of fibrosis, inflammation and copper associated genes in control vs. dietary mouse MASH mice post PBS or CuS administration. **(M)** – Schematic

### Chelation of Bioavailable Cu Reduces Fibrosis Independent of Hepatic Cu Restoration

Given the increase in circulating Cu observed in both genetic and dietary MASH models, and the inability of Cu supplementation to improve fibrosis, we hypothesized that excess bioavailable Cu contributes to stellate cell activation and fibrogenesis. To test this, mice subjected to a 4-week LMCD-HFD challenge were treated with the cell-impermeable Cu chelator bathocuproinedisulfonic acid (BCS) (Figure 8A).

**Figure 8.**
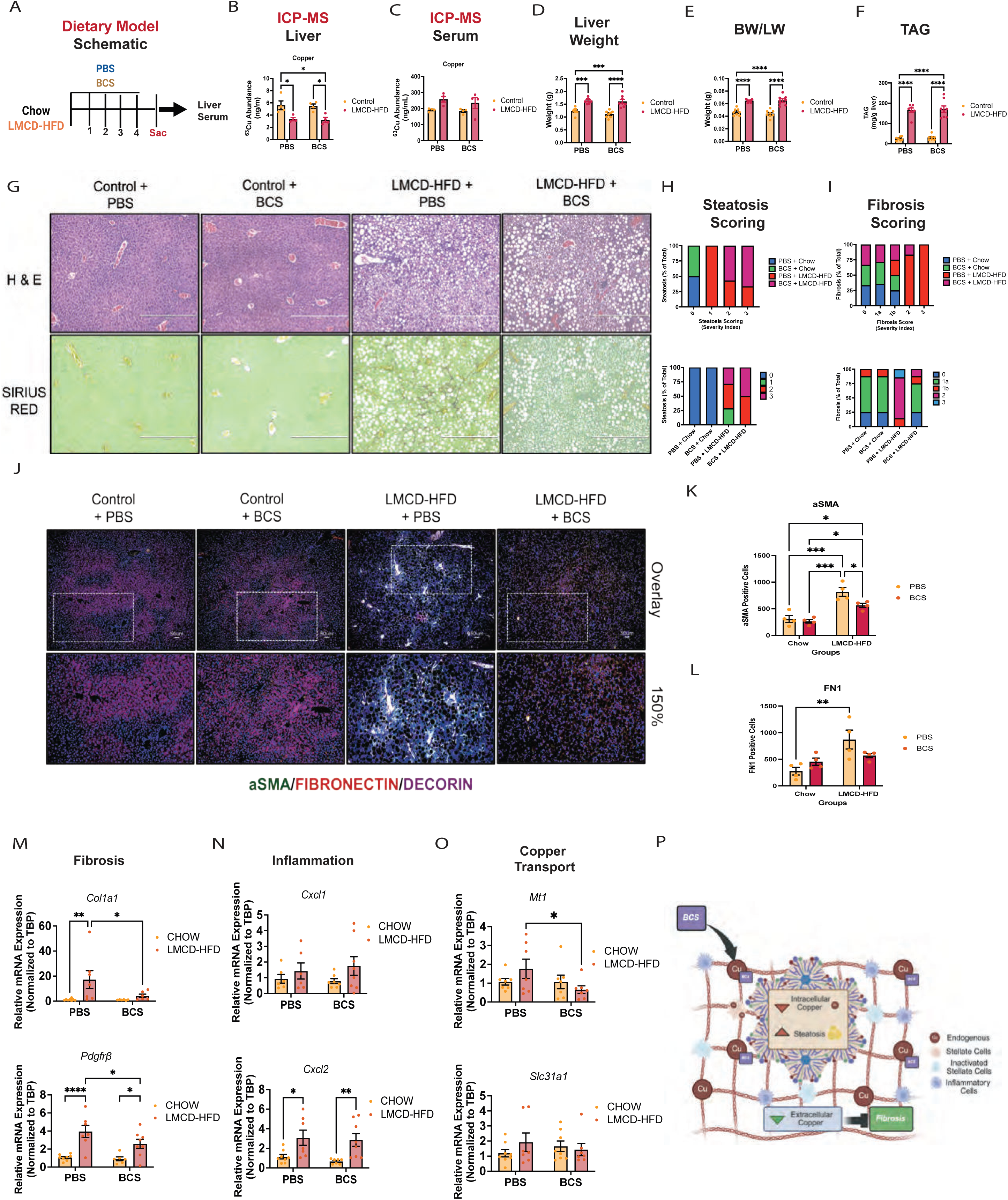
Chelation of Bioavailable Cu Reduces Fibrosis Independent of Hepatic Copper Restoration in MASH. **(A)** – Schematic of Study Design and Model. **(B)** – **(C)** – ICP-MS showing normalized ^63^Cu in the liver **(B)** or serum **(C)** of control vs. dietary MASH mice post PBS or 20 mg/kg BCS (daily/IP) for 4-wks. n = 7-8. **(D) - (E)** – Liver weight **(D)** and liver weight normalized to total body weight **(E)** after 4-wk challenge. **(F)** Triglyceride (TAG) assay quantifying liver fat accumulation in livers post PBS or BCS administration n = 7-8**. (G)** – Representative Histology. n = 7-8. **(H) – (I)** Blind fibrosis **(H)** and steatosis **(I)** scoring in control vs. dietary mouse MASH livers post PBS or BCS administration. n = 7-8. **(J)** – IF of αSMA and FN1-positive cells in of control vs. dietary MASH livers. **(K)** – **(L)** – Quantification of αSMA **(K)** and FN1-positive cells **(L)** in of control vs. dietary MASH livers. **(M)** – **(N)** - RT-qPCR of pro-fibrogenic **(M)** and pro-inflammatory **(N)** gene expression in control vs dietary mouse MASH livers +/- BCS. **(O)** – RT-qPCR of copper related gene expression in control vs dietary mouse MASH livers +/- BCS. Data are shown as mean ± SEM. *P < 0.05; **P < 0.01; ***P < 0.001. Scale bars: 400 μm.

BCS treatment did not restore hepatic Cu levels, as confirmed by ICP-MS (Figure 8, B and C), nor did it affect liver weight or hepatic triglyceride accumulation (Figure 8, D-F). Despite these minimal effects on hepatic lipid metabolism, BCS administration significantly reduced fibrosis, as assessed by Sirius Red staining (Figure 8G). Blinded histological scoring confirmed a shift toward lower fibrosis stages in BCS-treated mice, while steatosis severity remained unchanged (Figure 8, H and I).

Consistent with reduced fibrogenesis, BCS-treated livers exhibited decreased αSMA- and fibronectin (FN1)-positive cells (Figure 8, J-L), as well as reduced expression of pro-fibrogenic genes (Figure 8M). In contrast, inflammatory gene expression was not significantly altered (Figure 8N). Significant reduction in Cu genes *MT1A* and *ATP7B* were observed in BCS-treated livers (Figure 8O). Similar anti-fibrotic effects of BCS were observed in the genetic MASH model (Supplemental Figure 7). Together, these data demonstrate that Cu chelation selectively attenuates fibrosis without improving steatosis or hepatic Cu deficiency, supporting a role for compartment-specific Cu availability in regulating HSC-mediated fibrogenesis (Figure 8P).

### Combined Cu Supplementation and Chelation Do Not Provide Additive Anti-fibrotic Effects in MASH

Given that neither Cu supplementation (CuS) nor Cu chelation (BCS) alone normalized circulating Cu levels in MASH, we next tested whether combined treatment could restore Cu homeostasis and improve disease outcomes. Mice subjected to a 4-week LMCD-HFD challenge were treated with vehicle, BCS, CuS, or both agents (Dual Tx) (Figure 9A). As expected, CuS restored hepatic Cu content, whereas co-administration with BCS attenuated this effect (Figure 9, B and C). CuS treatment alone reduced liver weight and hepatic triglyceride accumulation and improved steatosis, while these effects were diminished in the Dual Tx group and absent in BCS-treated mice (Figure 9, D-H). Histological scoring confirmed that steatosis was most effectively reduced by CuS alone, with minimal improvement observed in Dual Tx mice (Figure 9J).

**Figure 9.**
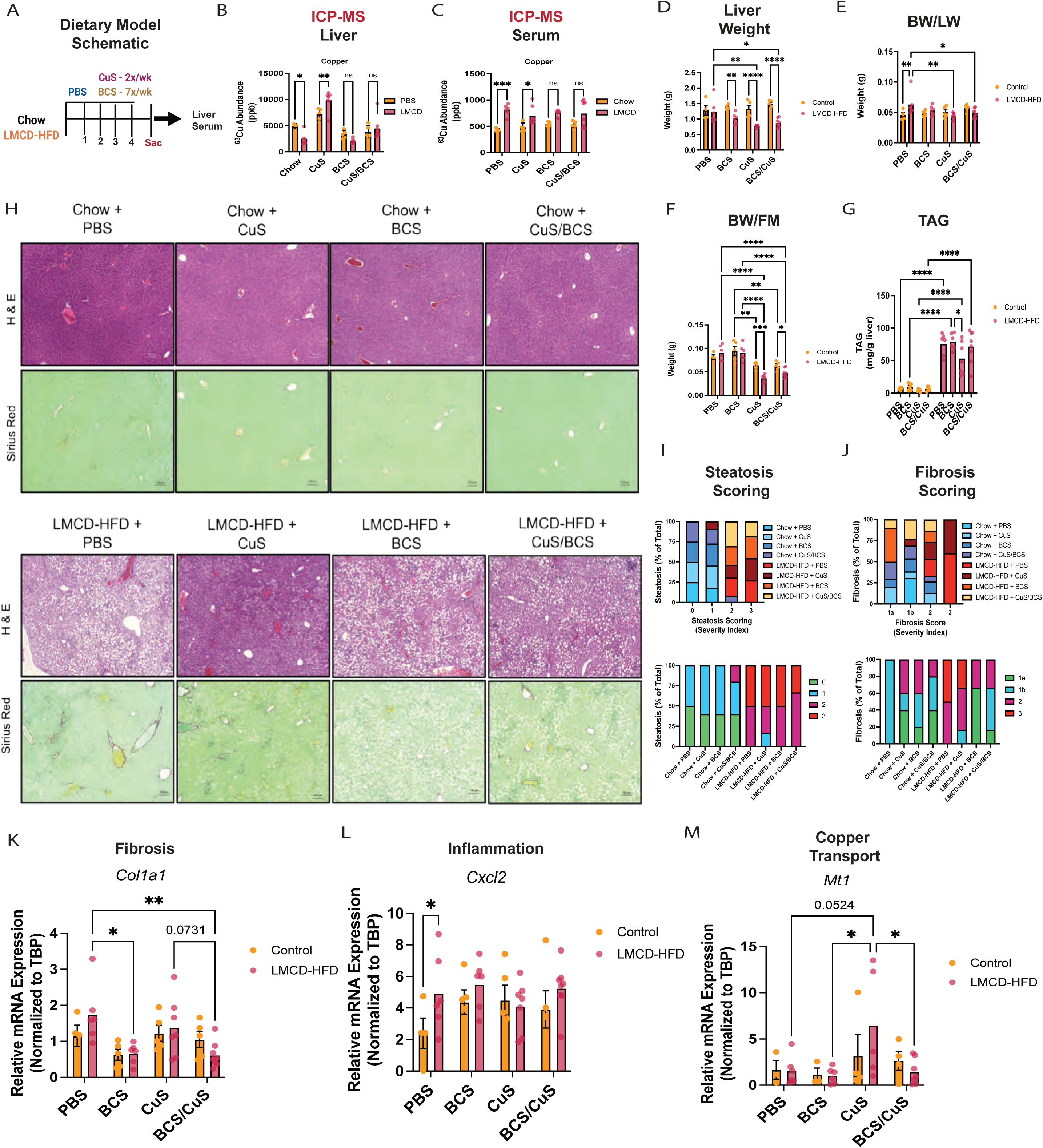
Combined Copper Supplementation and Chelation Do Not Enhance Anti-fibrotic Effects in MASH. **(A)** – Schematic of Study Design and Model. **(B)** – **(C)** – ICP-MS showing normalized ^63^Cu in the liver **(B)** or serum **(C)** of control vs. dietary MASH mice treated with either PBS (Vehicle), 20 mg/kg of cell-impermeable copper chelator - BCS (daily), 7.26 mg/kg of Gal-Cu (GTSM) (CuS) (2x/wk), or both BCS and CuS (Dual Tx) n = 5-6. **(D) - (E)** – Liver weight **(D)** and liver weight normalized to total body weight **(E)** after 4-wk challenge. **(F)** – Representative Histology. n = 5-6. **(G)** Fat mass normalized to BW in control vs. MASH mice 4-wks post drug administration. **(H)** Triglyceride (TAG) assay quantifying liver fat accumulation 4-wks post n = 5-6**. (I) – (J)** Blind steatosis **(I)** and fibrosis **(J)** scoring in control vs. dietary mouse MASH livers post drug administration. n = 5-6. **(K)** – **(L)** - RT-qPCR of pro-fibrogenic **(K)** and pro-inflammatory **(L)** gene expression in control vs dietary mouse MASH livers. **(M)** – RT-qPCR of copper related gene expression in control vs dietary mouse MASH livers. Data are shown as mean ± SEM. *P < 0.05; **P < 0.01; ***P < 0.001. Scale bars: 400 μm.

In contrast, fibrosis was significantly reduced by BCS treatment, with similar, though less pronounced, effects observed in Dual Tx mice (Figure 9I). Consistent with these findings, expression of pro-fibrogenic genes, including Col1a1, was reduced in both BCS- and Dual Tx-treated mice compared with CuS-treated and vehicle controls (Figure 9K). Inflammatory gene expression remained unchanged across all treatment groups (Figure 9L). *MT1A* expression, which was upregulated in response to CuS treatment, was significantly decreased in Dual Tx-mice (Figure 9M). Together, these data demonstrate that reduction of bioavailable Cu is required to mitigate fibrosis in MASH, whereas restoration of hepatocyte Cu is sufficient to improve steatosis. These findings highlight a functional dissociation between lipid accumulation and fibrogenesis driven by distinct, compartment-specific Cu pools.

## Discussion

Our study identifies disrupted Cu homeostasis as a consequence of impaired PC biosynthesis that contributes to MASH pathogenesis. We demonstrate that reduction of hepatic PC increases liver stiffness, impairs hepatocyte Cu uptake through mislocalization of the high-affinity Cu transporter CTR1 (*SLC31A1*), and results in decreased hepatic Cu content accompanied by increased circulating Cu. These findings define a mechanistic link between lipid remodeling and metal homeostasis in MASH.

Mechanistically, our data support a model in which impaired PC biosynthesis alters membrane composition and disrupts CTR1 localization at the plasma membrane, limiting hepatocyte Cu uptake. CTR1 is the primary high-affinity Cu importer in hepatocytes (76–78), and its mislocalization provides a direct explanation for the hepatic Cu deficiency observed in our models. As a result, hepatocytes cannot consume circulating Cu, increasing its availability to nonparenchymal compartments. *In vitro*, Cu directly promotes fibrogenic activation of HSCs, particularly in the presence of pro-fibrogenic stimuli such as TGFβ. Disruption of intracellular Cu import pathways attenuates fibrogenesis, indicating that intracellular Cu availability is required for HSC activation. Together, these findings support a model in which Cu functions as a signaling cofactor that amplifies fibrogenic responses.

HSC activation is a central driver of fibrosis regulated by mechanical cues, growth factor signaling, and ECM remodeling (5, 10, 18, 70, 74, 79–86). Our sNUC-RNA-Seq analyses reveal expansion of pro-fibrogenic HSC populations and enrichment of pathways associated with ECM organization and PDGFRβ and SMAD signaling, consistent with prior studies of HSC heterogeneity in MASH (86–89). Notably, we identified a quiescent HSC population enriched for Cu-associated genes in control livers, suggesting a role for Cu in maintaining HSC homeostasis, which disappears during disease progression (87). These findings align with previous work implicating ECM-associated pathways, including YAP/FAK signaling, matricellular proteins, and growth factor networks in HSC activation and fibrosis (28, 29, 90, 91). Cu-dependent lysyl oxidase (LOX) family members play a critical role in collagen crosslinking and extracellular matrix remodeling (5, 16–18, 92, 93). However, clinical targeting of LOXL2 with simtuzumab failed to improve liver or lung fibrosis (17, 92), suggesting that inhibition of individual Cu-dependent enzymes may be insufficient and supporting the need for strategies that more broadly modulate Cu availability across fibrogenic compartments.

Our data further suggest that Cu promotes fibrogenesis through MAPK-dependent signaling. Dysregulation of ERK/MAPK signaling is a well-established driver of HSC activation(98–102), and we observed Cu-dependent increases in ERK phosphorylation in stellate cells. Pharmacologic inhibition of ERK and p38 signaling attenuated Cu-induced fibrogenesis, consistent with prior work demonstrating that Cu can directly regulate kinase activity, including MEK1/2 in oncogenic signaling pathways(103). Although the precise molecular targets of Cu in HSCs remain to be defined, these findings support a broader role for Cu as a dynamic regulator of signaling networks involved in fibrosis.

Our findings also integrate with prior studies linking Cu dysregulation to liver disease, while providing new mechanistic insight. Both Cu deficiency and Cu overload have been associated with hepatic injury and fibrosis, highlighting the importance of tightly regulated Cu homeostasis(104–107).(108).(109).(84–90). For example, patients with Menkes disease, characterized by impaired Cu transport, exhibit hepatic injury and fibrosis (84–87), while Cu chelation has been shown to attenuate fibrosis in bile duct ligation models (88). Conversely, excessive Cu deficiency can exacerbate fibrosis (89), emphasizing the need for balanced Cu levels. Genetic models of Cu transport further illustrate this complexity: liver-specific *CTR1* deficiency does not recapitulate MASH phenotypes (77, 78, 91), whereas ATP7B deficiency leads to hepatocyte Cu accumulation and steatosis without fibrosis (92, 93), and global ATP7B loss results in severe inflammation and fibrosis (94–98). Together, these studies suggest that the pathological consequences of Cu dysregulation depend on its cellular and subcellular distribution.

Consistent with this concept, our *in vivo* data demonstrate that distinct pools of Cu differentially regulate disease features in MASH. Restoration of hepatocyte Cu using a liver-directed Cu ionophore improved steatosis but failed to reduce fibrosis, whereas pharmacologic Cu chelation attenuated fibrosis without correcting hepatic Cu deficiency. Moreover, combined Cu supplementation and chelation did not provide additive benefit over chelation alone. These findings demonstrate a functional dissociation between lipid accumulation and fibrogenesis and support a model in which compartment-specific Cu availability, rather than total Cu levels, governs disease progression. In this framework, hepatocyte Cu primarily regulates lipid metabolism, whereas bioavailable Cu accessible to HSCs promotes fibrogenesis.

It is important to note however, that dysregulations in essential metals other than Cu, also occur in MASH mouse livers and serum. Similar to the trend observed with Cu, MASH livers also showed significant decreases in hepatic selenium and increases in serum selenium (Supplemental Figure 8, A and B). Interestingly, patients with MASH report hepatic selenium deficiency, with decreases in selenoproteins (selenium binding proteins) leading to impairments in PPARα-mediated fatty acid oxidation (FAO) (106). Selenium supplementation is currently under investigation as a therapeutic against MASH (107). MASH mice also demonstrated significant increases in serum iron (trending increase in hepatic iron) (Supplemental Figure 8B). ∼40-60% of MASH patients report increases in serum iron and ferritin (iron binding protein) (108). Hepatic iron overload also associates with ferroptosis, (iron-mediated cell death), disease severity (advanced fibrosis), and mortality in MASLD and MASH patients (108–110). Research into the use of iron chelators, such as deferoxamine, as a therapeutic against MASH is ongoing (109, 111–113). Lastly our MASH models showed a significant decrease in serum zinc levels (Supplemental Figure 8B). Zinc deficiency is prevalent in MASLD and MASH patients, with over 80% of those with advanced fibrosis (cirrhosis) affected (114–118). Treatment with PPARα agonist – Pemafibrate, effectively restores zinc levels and mitigates fibrosis in biopsy-confirmed MASH patients (119). Moreover, studies have demonstrated the benefits of direct zinc supplementation through improvements in diet-induced hepatic steatosis, inflammation, and liver injury (120–123), findings also recapitulated in patients with chronic liver diseases such as MASLD/MASH (124–128). These data suggest that hepatic lipid remodeling in MASH may also affect the homeostasis of other essential metals. Future studies need to investigate the pathogenic or hepatoprotective roles of these metals in MASH.

Emerging evidence suggests that phospholipids play an important role in regulating Cu homeostasis and transporter function. In non-mammalian systems, phosphatidic acid can inhibit CTR1 activity (129), and reconstitution studies indicate that a phosphatidylcholine-rich membrane environment is required for proper CTR1 function (130–133). These findings support a model in which membrane lipid composition directly influences Cu transport. Consistent with this, our data demonstrate that acute reduction of PC biosynthesis is sufficient to impair CTR1 localization at the plasma membrane, providing direct *in vivo* evidence that PC-dependent membrane remodeling regulates hepatocyte Cu uptake.

Functionally, our *in vivo* studies reveal that restoration of hepatocyte Cu improves steatosis but does not resolve fibrosis, whereas Cu chelation attenuates fibrogenesis without correcting hepatic Cu deficiency. Moreover, combined Cu supplementation and chelation do not provide additive benefit, further supporting the concept that distinct Cu pools differentially regulate disease features. These findings suggest that targeting overall Cu levels may be insufficient and instead highlight the importance of modulating site-specific Cu availability. Consistent with this idea, prior studies have shown that liver-directed Cu supplementation can improve lipid metabolism in MASLD(79, 138, 139), while Cu chelators such as tetrathiomolybdate have demonstrated anti-fibrotic effects in other disease contexts (88).

Together, these observations support a model in which PC-dependent membrane organization governs Cu distribution, linking lipid metabolism to fibrogenic signaling. These data further suggest that therapeutic strategies aimed at selectively restoring hepatocyte Cu while limiting bioavailable Cu in fibrogenic compartments may represent a novel approach for treating MASH. Future studies will be required to define the precise molecular mechanisms by which Cu regulates stellate cell activation and to identify strategies for selectively targeting these pathways.

## Methods

### Sex as a biological variable

This study exclusively examined male mice; however, we expect similar results in female mice, due to the well-established role of hepatic phosphatidylcholine deficiency, in the onset of MASH-associated fibrosis and inflammation, independent of sex (36, 65).

### Animal Experiments

8-10-wk old, C57BL/6J mice were obtained from the Jackson Laboratory (Catalog # 000664), and housed, under specific pathogen-free conditions in facilities at the University of Pennsylvania. Mice were housed in the animal facility with a 12-hour dark/light cycle (7:00 a.m. to 7:00 p.m.) and *ad libitum* access to food and water. For acute excision of liver-specific genes, mice were injected retro-orbitally with AAV (Vector Core, University of Pennsylvania) containing a liver-specific thyroxine binding globulin (TBG) promoter serotype 8 (AAV8-TBG) containing either GFP (AAV-GFP) or Cre (AAV-Cre) at a dosage of 1.0 × 10^11^ genome copies. A subset of mice were fed a chow diet (LabDiet, no. 5010), whereas the dietary MASH mice were fed a LMCD-HFD (45 kcal% fat with 0.1% methionine; no added choline) for 4-wks (Research Diets, #A06071309), as previously described (58). A subset of mice (control) were administered 1X PBS, whereas other mouse subsets were intraperitoneally injected with bathocuproinedisulfonic acid, BCS (20 mg/kg) (Sigma-Aldrich, B1125) daily for one month as previously described (105, 137). All experiments were performed in male mice. 7.26 mg/kg of Cu supplement [CuS; Gal-Cu (GTSM)] Cu supplement (CuS) was administered twice per week, via intraperitoneal injection as previously reported (135). Liver pathologies were evaluated blindly, by independent scorers as per this published rubric (138).

### Whole-Body Metabolic Analysis

Mice had free access to food and water. Body fat composition (lean mass; fat mass) was assessed in ad-libitum fed mice, by nuclear magnetic resonance imaging (ECHO MRI-Rodent Metabolic Phenotyping core (RMPC) mouse, University of Pennsylvania-Perelman School of Medicine).

### Nuclei Isolation

Nuclei were isolated from frozen liver samples (∼200 mg/ 0.20 g) as previously described (dx.doi.org/10.17504/protocols.io.3fkgjkw) (139). Livers were weighed out on dry ice and homogenized in Lysis Buffer (see Table 1) using a large glass dounce homogenizer. Homogenates were filtered using a 70-μm cell strainer, transferred to microcentrifuge tube, and centrifuged at 500 *g* and 4°C for 5 minutes. The supernatant was removed, and fresh Lysis Buffer was added for an additional 5 minutes on ice followed by centrifuged at 500 *g* and 4°C for 5 minutes. Trypan blue was used to monitor nuclei morphology between isolation steps. Nuclei pellet (>700-1,200 nuclei/μl), was washed twice in Homogenization buffer (see Table 2), filtered through a 40-μm cell strainer, and immediately processed for single nuclei RNA-Seq at the Center for Applied Genomics Core at Children’s Hospital of Philadelphia.

### Single-Nuclei RNA Sequencing Data Analysis

Libraries were prepared using the 10x Genomics Chromium Single Cell 3′ v3 kit. Sequencing was performed using an Illumina NovaSeq 6000 per manufacturer’s instructions. After sequencing, Cell Ranger v3.0.2 (10x Genomics) was used to align sequencing reads to a custom reference genome (mouse mm10 release 93 genome build), which included introns and exons to account for pre-mRNA and mature mRNA present in the nucleus. Raw counts were further analyzed using both Seurat v3.1.1, and v5(143, 144). Each sample was filtered for (1) genes expressed in at least 3 nuclei, (2) nuclei that express at least 100 genes, and (3) ≤1% mitochondrial genes. Additional quality control was performed using the scatter package (v1.10.1). The DoubletFinder v2.0.2 package excluded putative doublets from subsequent analyses. For gene-level analysis, ‘counted’ RNA molecules are redefined as UMIs (unique molecular identifiers), to account for PCR amplification bias in NGS reads. UMI counts are then normalized as an aggregated object (containing all 4 libraries), Feature counts (UMI counts) for each cell are taken and divided by the total counts for that cell and multiplied by a scale factor (where the scale factor is scale.factor = 1e6). Values are then natural log transformed using log1p. ScaleData was used to perform feature (gene)-level scaling, meaning that each feature was centered at 0 and scaled by its standard deviation. This normalization and scaling of all cells within each group allow for sample-level and group comparisons. To perform sample integration, a Harmony-based workflow was used, an algorithm that corrects for “batch effects” between samples (https://github.com/immunogenomics/harmony) (145). Downstream PCA, clustering, and UMAP projections were performed using Seurat within R. Differentially expressed genes were determined using a Wilcoxon-rank sum test with a Bonferroni correction to account for multiple comparisons.

### Human Single-Cell RNA Sequencing

These data was obtained through previously published publicly available work available here(146). Normal human livers and NASH human livers used for single-cell RNA sequencing were obtained through the Liver Tissue Cell Distribution System, Minneapolis, Minnesota, which was funded by National Institutes of Health contract no. HSN276201200017C.

### ICP-MS Analysis

Frozen tissues were homogenized mechanically using a mortar and pestle system over dry ice. A portion of the homogenized tissue was collected and weighed. The tissue was then digested overnight using concentrated trace metal-grade nitric acid (Fisher, UN2031) at a volume of 2 µL per 1 mg tissue. The following day, the digest was diluted 25× with ultrapure water and centrifuged at maximum speed for 30 minutes to pellet debris. The supernatant was collected and the centrifugation repeated to fully clarify the sample. Serum samples were prepared by diluting 10 µL of thawed serum 25× with a solution of 4% nitric acid in ultrapure water. Samples were then immediately centrifuged at maximum speed for 30 minutes, and the supernatant was collected.

All samples were measured by ICP-MS (iCAP TQ, Thermo Fisher Scientific) using the helium kinetic energy discrimination (KED) mode to reduce oxide interference. A standard curve of 0, 10, 100, 500, and 1000 ppb analytes was made by diluting Inorganic Ventures Stock Solutions CMS-4 and CMS-5 with 4% nitric acid in ultrapure water. A stock solution of 100 ppb Germanium-73 was used as an internal standard.

### LA ICP-MS Analysis

For each frozen liver tissue, 25μm thick sections were sectioned on a cryostat (Leica, CM1850). Sections were thaw-mounted on poly-L-lysine coated glass slides (Sigma Aldrich) and desiccated under vacuum for 10min. The LA-ICP-MS imaging was carried out at 15μm resolution with the Iridia Laser Ablation Unit (Teledyne Technologies) coupled with iCAP TQ ICPMS (Thermo Fisher Scientific) instrument. The laser was set at a rep rate of 500Hz, and fluence was set to 1J/cm2. The image was processed using HDIP Mass Spectrometry Data Analysis software (Teledyne Technologies).

### Histology and IHC-based staining

For tissue-based or immunohistochemistry (IHC), mouse livers were dissected and fixed in 10% formalin. Following tissue embedding into paraffin blocks, embedded tissues were cut into 10 μM-thick sections placed on SuperFrost slides (Thermo Scientific). Following de-paraffinization, and antigen retrieval, slides were then stained with: Hematoxylin and Eosin (H&E) or Picro Sirius Red Stain Kit (Connective Tissue Stain), or Primary antibodies against alpha Smooth Muscle Actin (Thermo # 53-9760-82 - 1:200), CoraLite594-conjugated Fibronectin (ProteinTech # CL594-66042 (1:200), Decorin (Thermo # 14667-1-AP - 1:200), and Cytoglobin (Thermo # 60228-1 -1:200) according to manufacturers’ protocols.

### Immunofluorescence

For cell-based immunofluorescence (IF), samples were fixed with 10% paraformaldehyde and then blocked for 30-min at room temperature using 10% FBS/0/2% Triton-X-100/1X PBS. The following primary antibodies were then used to stain respective cell lines: Pre-conjugated *SLC31A1*/CTR1 antibody (NOVUS BIO no. 402AF594), ATP7B antibody (Proteintech (no. 19786-1-AP), both at a concentration of 1:200, and Pre-conjugated Fibronectin antibody (Proteintech no. CL594-15613) at a concentration of 1:1,1000. Alexa Fluor secondary antibodies from Invitrogen Life Technologies, were used. Hoescht was used for nuclear counterstaining.

### CAP-1 Cu +1 Probe Imaging

For live-cell based imaging of Cu uptake we utilized a Cu^+1^ selective probe (CAP-1), that emits fluorescence at 488 nm in the presence of reduced Cu (excitation wavelength: 488 nm, emission wavelength 500-600 nm). In brief, following treatment with either: serum free media (SFM), 500 μM Cu chelator BCS, or 100uM CuSO_4_ for 6-hr, cells were washed using 1x PBS, then incubated with serum-free media containing 1 uM CAP-1 for 40 minutes at 37 degrees Celsius. Cells were then washed with pre-heated 1x PBS (3 x 5 min), then incubated for 5 min at 37 degrees Celsius with 1x PBS containing 1 uM CAP-1. Cells were subsequently washed with 1x PBS (3 x 10 min), then imaged using the EVOS 7000 Fluorescence microscope.

### Immunoblots

Protein lysates were prepared from frozen livers in a modified RIPA buffer with Phosphatase Inhibitor Cocktails 2 and 3 (Sigma-Aldrich) and Complete Protease Inhibitor Cocktail, as previously described(60, 62, 147). The following antibodies were used for immunoblotting, all from Cell Signaling Technology: HSP90 at 1:1,000 (CST no. 4874), phsopho-p44/42 ERK1/2 (Thr202/Tyr204) at 1:1,000 (CST no. 76939S), total-ERK1/2 at 1:1,000 (CST no. 4695S), and CTR1 at 1:1,000 (CST no. 13086S). The following antibody was used for immunoblotting from ProteinTech: ATP7B at 1:10,000 (no. 19786-1-AP). The following antibody was used for immunoblotting from Santa Cruz: Smooth Muscle Actin (B4) at 1:1,000 (SC no. 53142). 5% dry milk in TBST, or Intercept Blocking Buffer (LICOR no. 927-60001) was used to block nitrocellulose membrane (BioRed # 1620115), following TurboTransfer.

### mRNA isolation and real-time PCR

Total RNA was isolated from frozen livers using the RNeasy Plus kit (Qiagen). Complementary DNA was synthesized using Moloney murine leukemia virus (MulV) reverse transcriptase, and the relative expression of the genes of interest was quantified by real-time PCR using the SYBR Green dye-based assay.

### Histology

Livers were fixed in 10% buffered formalin overnight, dehydrated in ethanol, paraffin-embedded, and sectioned. Sections were stained with H&E or Sirius red staining.

### Statistics

Statistical analysis was performed using 1-way ANOVAs when more than 2 groups were compared, 2-way ANOVAs when 2 conditions were analyzed, followed by Sidak’s multiple comparisons test (2 groups), or Tukey multiple comparisons test (>2 groups), and unpaired 2-tailed Student *t* test when 2 groups were being assayed. All data were presented as mean ± SEM. *P value < 0.05, **P value < 0.01, ***P value < 0.001, ****P value < 0.0001 versus indicated treatment groups.

### Study Approval

The institutional animal care and use committees of the University of Pennsylvania approved all the animal studies, which adhered to the National Institutes of Health guidelines for the care and use of laboratory animals.

## Data Availability

All single nuclei RNA sequencing data are available in NCBI SRA (Sequence Read Archive) by the following accession numbers: GSE330606. Values for all data points in graphs are reported in the Supporting Data Values files. Materials such as cell lines are available from primary and corresponding authors.

## Supporting Data Files

Figure 2F – Gene Enrichment List – Top 100 DEG Pathway Analysis – Genetic MASH Liver – Total

Figure 2G – Gene Enrichment List - Top 100 DEG Pathway Analysis – Dietary MASH Liver - Total

Figure 2H – Gene Enrichment List - Top 100 DEG Pathway Analysis – Shared MASH Liver – Total

Figure 3D – Gene Enrichment List - Pathway Analysis – Genetic MASH Liver – Hepatocyte cells

Figure 3E – Gene Enrichment List - Pathway Analysis – Dietary MASH Liver – Hepatocyte cells

Figure 3F – Gene Enrichment List - Pathway Analysis – Human MASH Liver – Hepatocyte cells

Figure 4B – Gene Enrichment List – Volcano Plot – Genetic MASH Liver – Stellate cells

Figure 4C – Gene Enrichment List – Volcano Plot – Dietary MASH Liver – Stellate cells

Figure 6C – Gene Enrichment List - STRING Pathway Analysis – Dietary MASH Liver – Stellate cells

Figure 6D – Gene Enrichment List - STRING Pathway Analysis – Genetic MASH Liver – Stellate cells

Supplemental Figure 2B – Gene list for DEGs Heatmap - Control vs. Genetic and Dietary MASH Liver – Total

Supplemental Figure 2C – Gene list for Venn Diagram – DEGs in Genetic/Dietary MASH Liver – Total

## Acknowledgments

Lan Cheng - for expert tissue processing and histology

Clementina Messaros and Jimmy Xu – Biomolecular Mass Spectrometry Core, Center of Excellence in Environmental Toxicology for ICP-MS analysis of mouse serum and livers

Rene Jacobs – for generation and distribution of L-*PCYT1A* - *fl/fl* mice

Dylan Weissenkampen – for consult on single nuclei analysis tools

Joe Baur – for guidance and study support

## Author contributions

J.W. conceived the hypothesis, designed, and performed experiments, analyzed data, and prepared the manuscript. M.V.B. and J.P.G. analyzed transcriptomics data. D.L. and R.G.W. performed rheology experiments and analyzed rheometric data. D.A.S. performed blind steatosis and fibrosis scoring. G.M.N, F.K.C., C.J., and R.W.M performed ICP-MS and Spatial ICP-MS. Y.X., J.Z., and M.A.L. performed and analyzed human MASH single-cell RNA sequencing experiments. G.L. provided technical assistance with Cu probe imaging. J.K., Y.A., and C.J.C. provided technical assistance and reagents for Cu supplementation studies. D.C.B. helped design the experiments, prepare the manuscript, and co-directed the project. P.M.T. conceived the hypothesis, designed experiments, helped prepare the manuscript, and directed the project.

## Abbreviations

ALT, AST, BCS, CCTα, CHO, CRP, Cu, CuS, ERK1/2, Fe, Gal-Cu, GAN, HEP, HFD, HSC, KC, LMCD-HFD, LSEC, MAPK, MASLD, MASH, MEK1/2, PC, PEMT, scRNA-seq, Se, snRNA-seq, vLDL, Zn

