## Supplementa Figures 1-8, Tables 1 & 2 for "Reduction in Hepatic Phosphatidylcholine Biosynthesis Promotes MASH Through Copper Deficiency"

#### Supplemental Figure 1

**Reduced Hepatic PC Biosynthesis Induces MASH-associated Gene Expression Markers in MASH Livers**

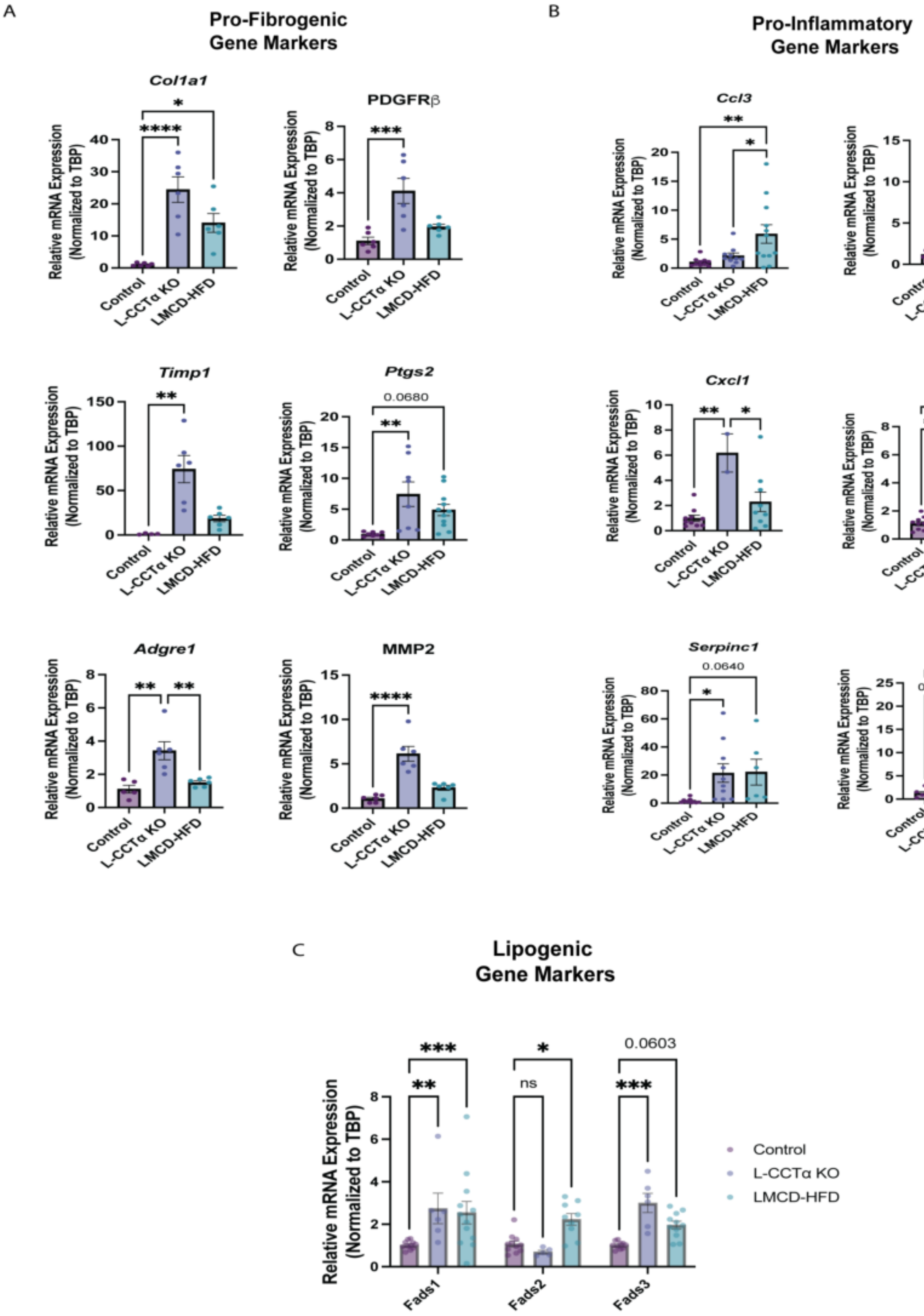


**(A)** RT-qPCR showing induction of pro-fibrogenic gene expression with increased stiffness in mouse MASH livers compared to control. n = 6. **(B)** RT-qPCR showing induction of pro-inflammatory gene expression with increased stiffness in mouse MASH livers compared to control. n = 6. (**C)** RT-qPCR showing change in lipogenic gene marker - FADS1-3 in mouse MASH livers compared to control. n = 6. Data are shown as mean ± SEM. *P < 0.05; **P < 0.01; ***P < 0.001

#### Supplemental Figure 2

##### Hepatic Cell Populations and Zonation Patterns Are Defined in Mouse MASH Models

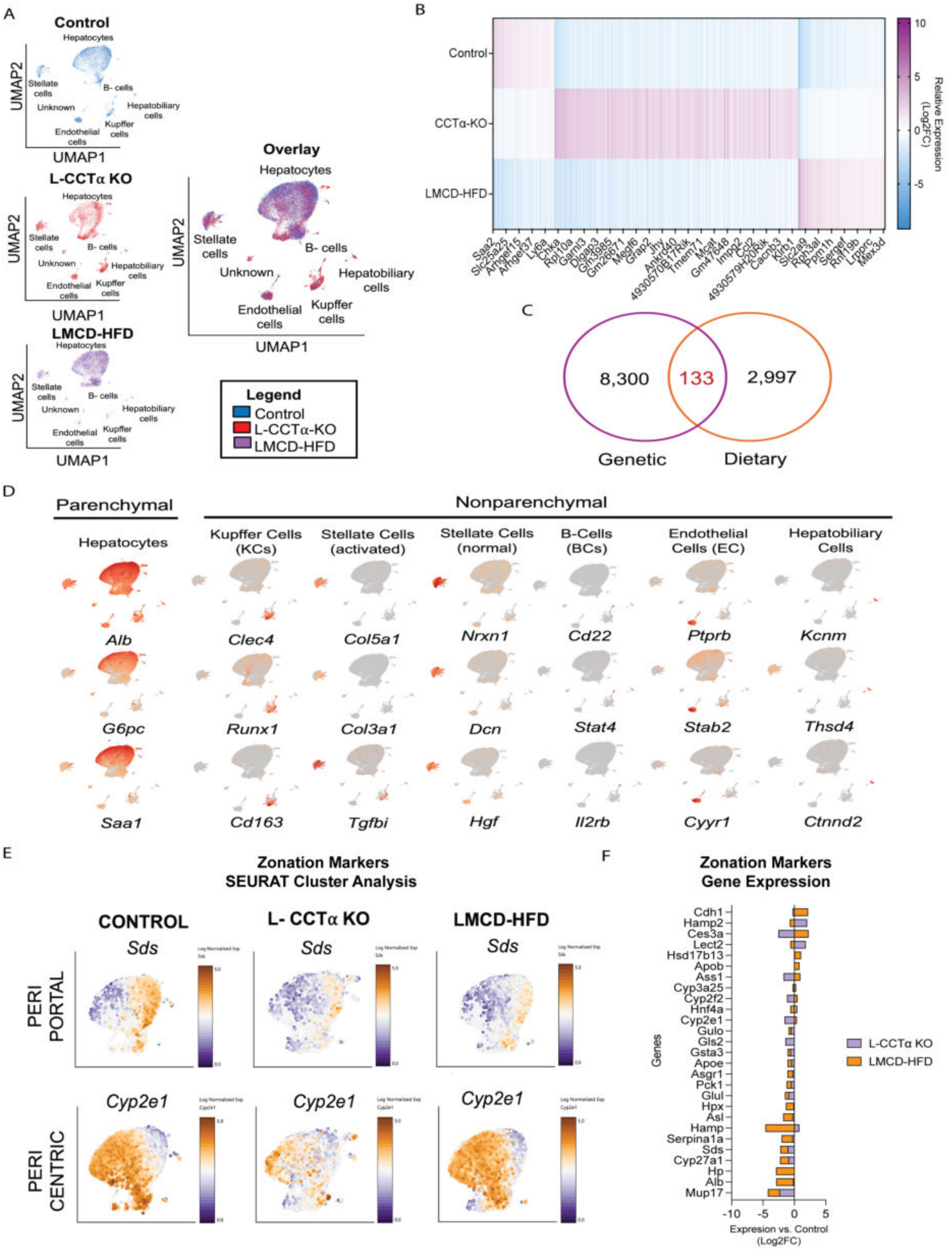


**(A)** – Populations of nuclei clustered and color-coded by group between mouse MASH and control livers. **(B)** Heat map of DEGs identified in total Control vs Mouse MASH livers (sorted by p-value <0.05). **(C)** – Venn diagram of DEGs identified in total liver and shared between both mouse MASH livers compared to control. **(D)** – Annotation of specific clusters identified using previously established human/mouse cell markers. **(E)** – SEURAT Cluster heat map showing periportal and pericentral zonation marker expression. **(F)** – Graph illustration Log2FC of Zonation-associated gene expression markers in genetic and dietary MASH livers.

### Supplemental Figure 3

#### Copper Homeostasis-Related Gene Expression Is Dysregulated in Mouse MASH Hepatocytes

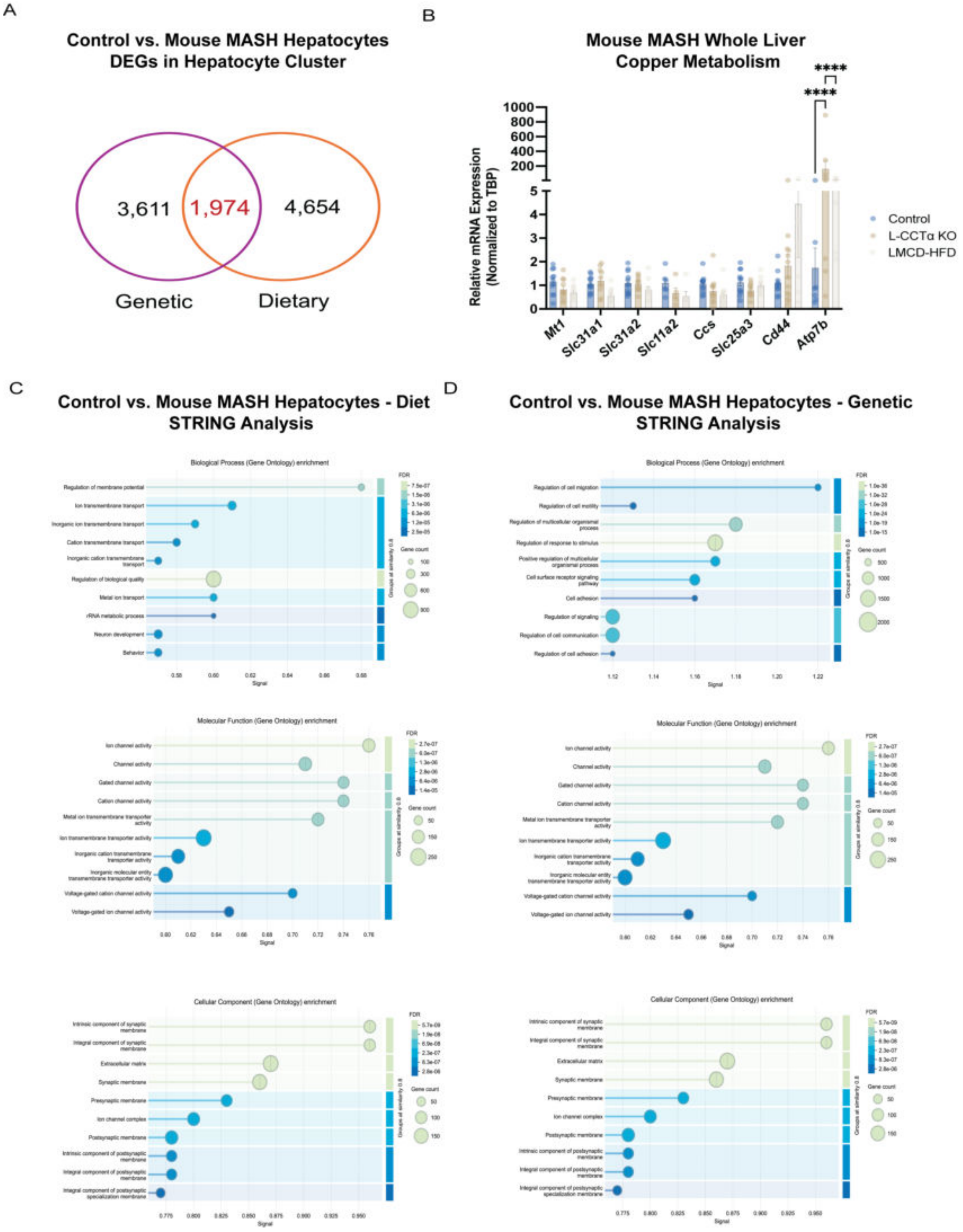


**(A)** - Venn Diagram of DEGs shared between genetic and dietary mouse MASH-derived hepatocytes. **(B)** RT-qPCR showing the expression of copper associated signaling genes from genetic and dietary MASH livers compared to control. n = 5-6. **(C)** –STRING Cluster analysis for Biological Process (Gene Ontology), Molecular Function (Gene Ontology), and Cellular Component (Gene Ontology) analysis for dietary, and **(D)** – genetic mouse MASH hepatocytes. False discovery rates illustrated in dot plot, with gene # demonstrated by size difference. Data are shown as mean ± SEM. ****P < 0.0001; P > 0.05

### Supplemental Figure 4

#### Loss of Quiescent and Expansion of Activated Hepatic Stellate Cells in MASH

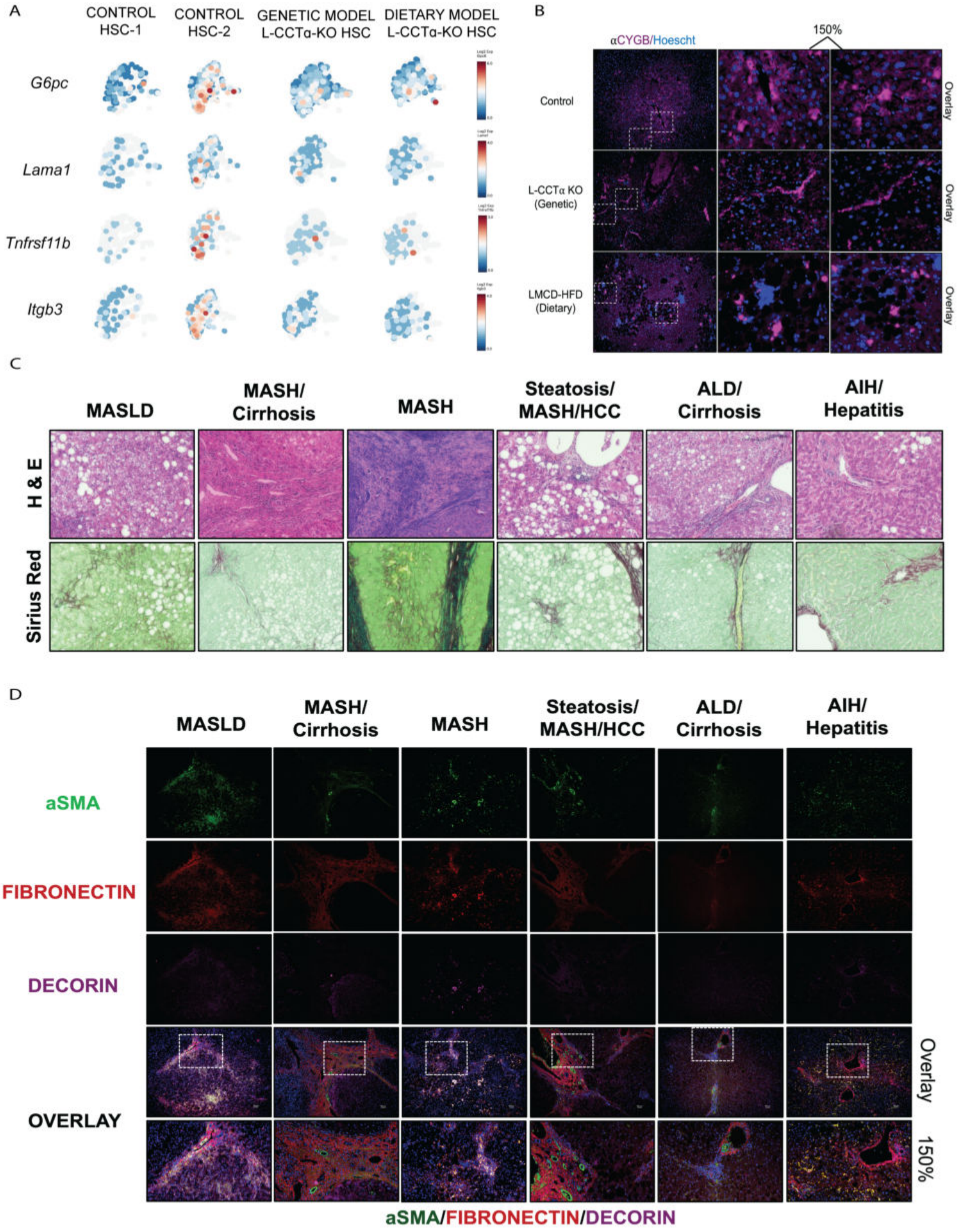


**(A)** U-Maps showing gene expression overlay of quiescent genes in control vs. mouse MASH HSCs. **(B)** IF stain of quiescence HSC marker – CYGB, in control vs. mouse MASH livers. n = 3 Explain what the second. **(C)** – Representative histology (H&E and Sirius Red) of human liver diseases. **(D)** – IF stain of human liver sections probing for activated HSCs using antibodies against α-SMA, Fibronectin, and Decorin. Scale bars: 400 μm.

### Supplemental Figure 5

#### PC Deficiency Disrupts CTR1 Localization and Impairs Hepatocyte Copper Uptake

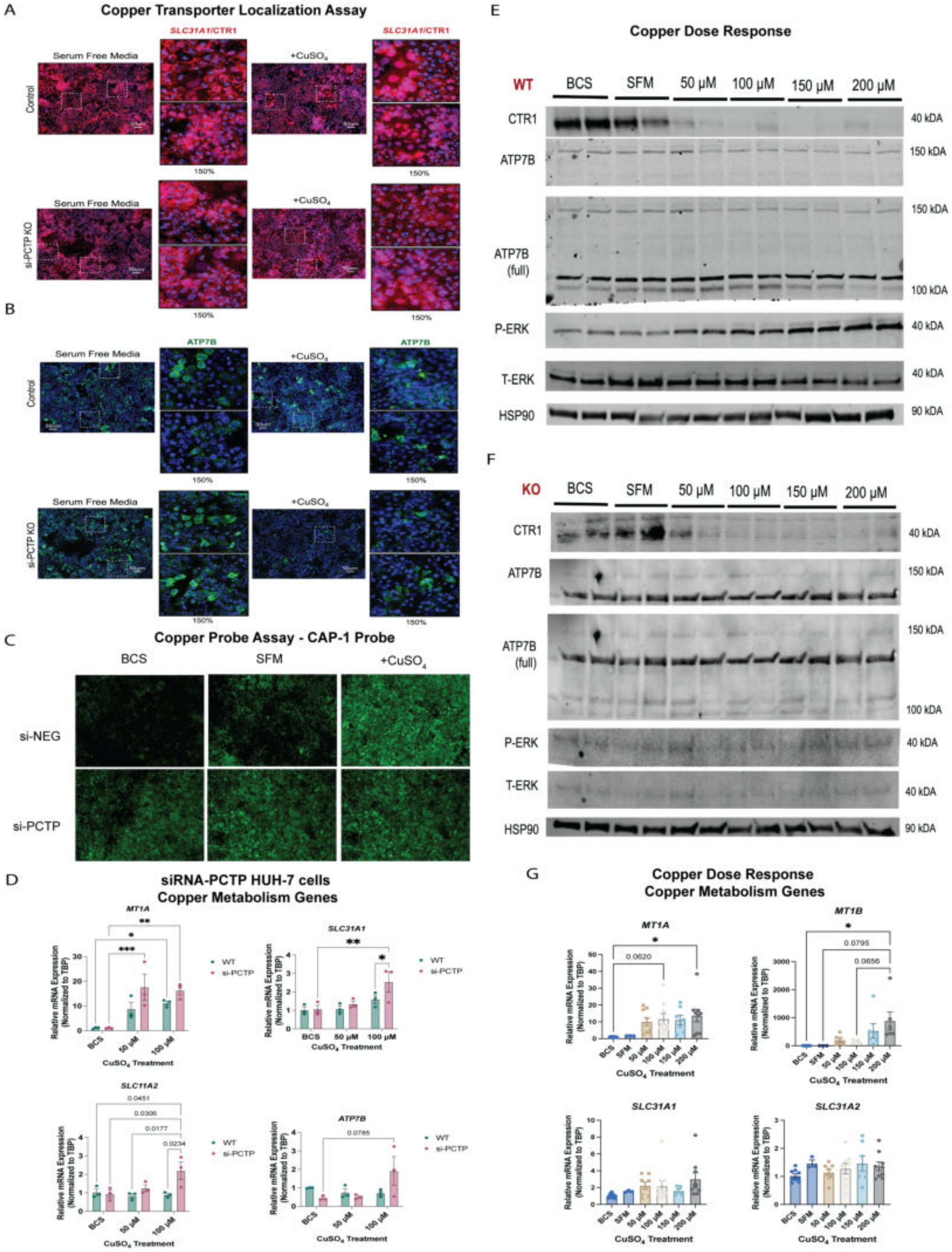


IF showing CTR1 **(A)** and ATP7B **(B)** localization in siRNA-negative control or siRNA-*PCTP* treated HUH-7 cells. Wildtype control cells were from the same experiment as Figure 5A. n = 3-4. **(C)** – IF showing Cu import using CAP-488, a $Cu^{+1}$ specific probe that emits a fluorescence at 488 in the presence of copper in HUH-7 cells transfected with siRNA against *PCTP.* n = 3-4. **(D)** RT-qPCR of HUH-7 cells in the presence or absence of either siRNA-Negative Control or siRNA-*PCTP* after being treated with either Serum Free Media (SFM), 500 µM BCS, 50 µM, or 100 µM $CuSO_4$ for 6-hr. n = 3 per group. **(E)** – Western blot analysis of wildtype HUH-7 cells transfected with siRNA-negative control and treated with an increasing dose of $CuSO_4$ for 6-hr. **(F)** – Western blot analysis of HUH-7 cells transfected with siRNA-*PCYT1A* for 48-hr, then treated with an increasing dose of $CuSO_4$ for 6-hr. **(G)** – RT-qPCR analysis of HUH-7 cells subjected to a 6-hr $CuSO_4$ dose response. n = 3-8. Scale bars: 50 µm. Data are shown as mean ± SEM. $^*P < 0.05$; $^{**}P < 0.01$; $^{***}P < 0.001$; NS, no significance, $P > 0.05$

### Supplemental Figure 6

#### Copper Promotes Fibrogenic Activation of Hepatic Stellate Cells in a MAPK-Dependent Manner

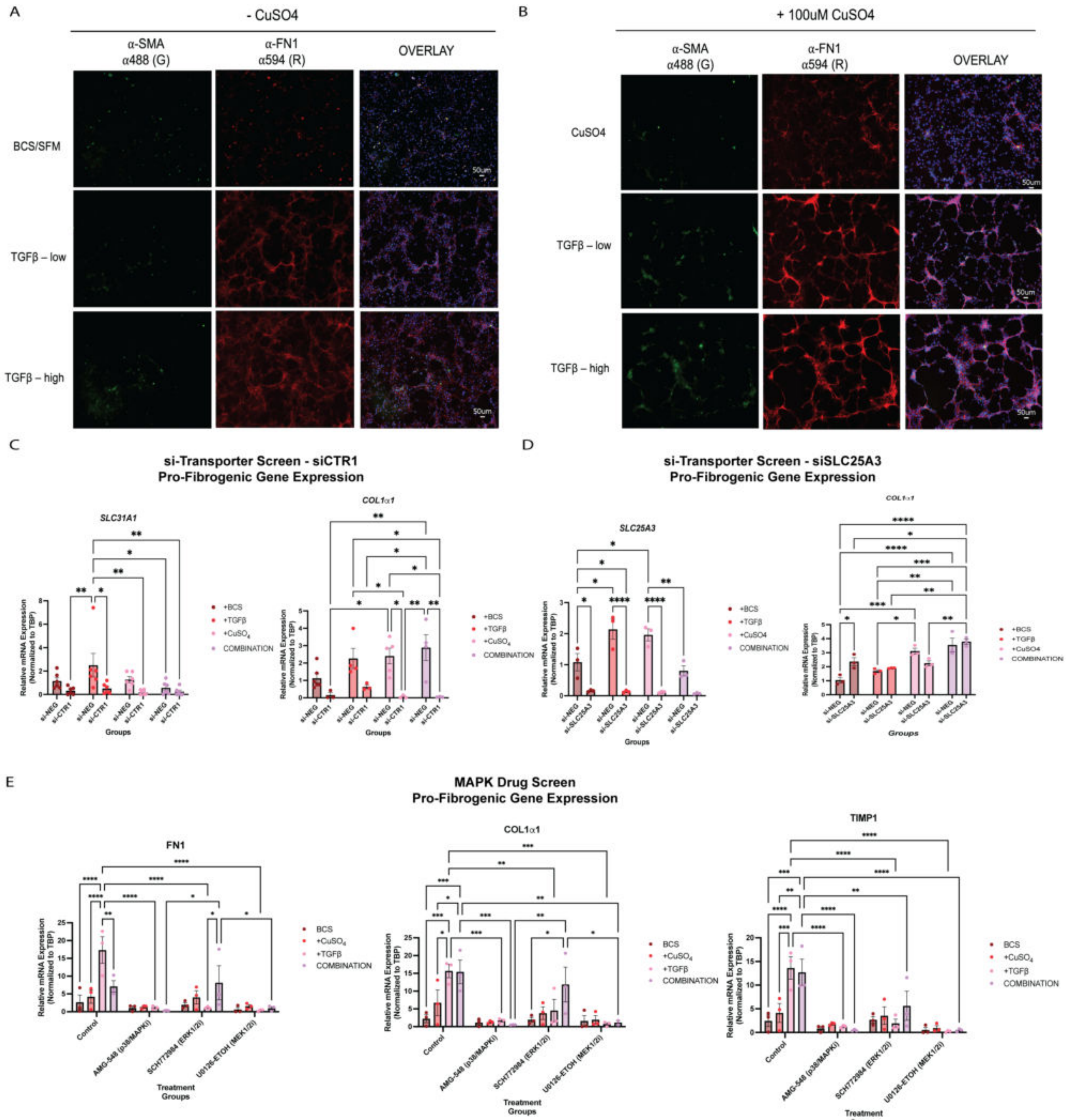


**(A) – (B)** - IF showing ECM protein – Fibronectin (FN1) expression and matrices formation increases with both $CuSO_4$ and low – 10 ng/mL, or high – 100 ng/mL TGFB treatment for 24-hrs. Scale bars: 50 µm. **(C)** RT-qPCR showing pro-fibrogenic gene expression in CTR1 knockdown cells. n = 5-6. **(D)** – RT-qPCR showing pro-fibrogenic gene expression in SLC25A3 knockdown cells. n = 3-4. **(E) –** RT-qPCR showing pro-fibrogenic gene expression in cells pre-treated for 1-hr with MAPK inhibitors – AMG-548 (p3/MAPK), SCH772584 (ERK1/2), or U0126 (MEK1/2), then treated with $CuSO_4$ and/or 10 ng/mL TGF$\beta$. n = 3-6. Data are shown as mean ± SEM. *P < 0.05; **P < 0.01; ***P < 0.001; NS, no significance, P > 0.05. Scale bars: 400 µm.

### Supplemental Figure 7

#### Copper Chelation Reduces Fibrosis in Genetic MASH Models

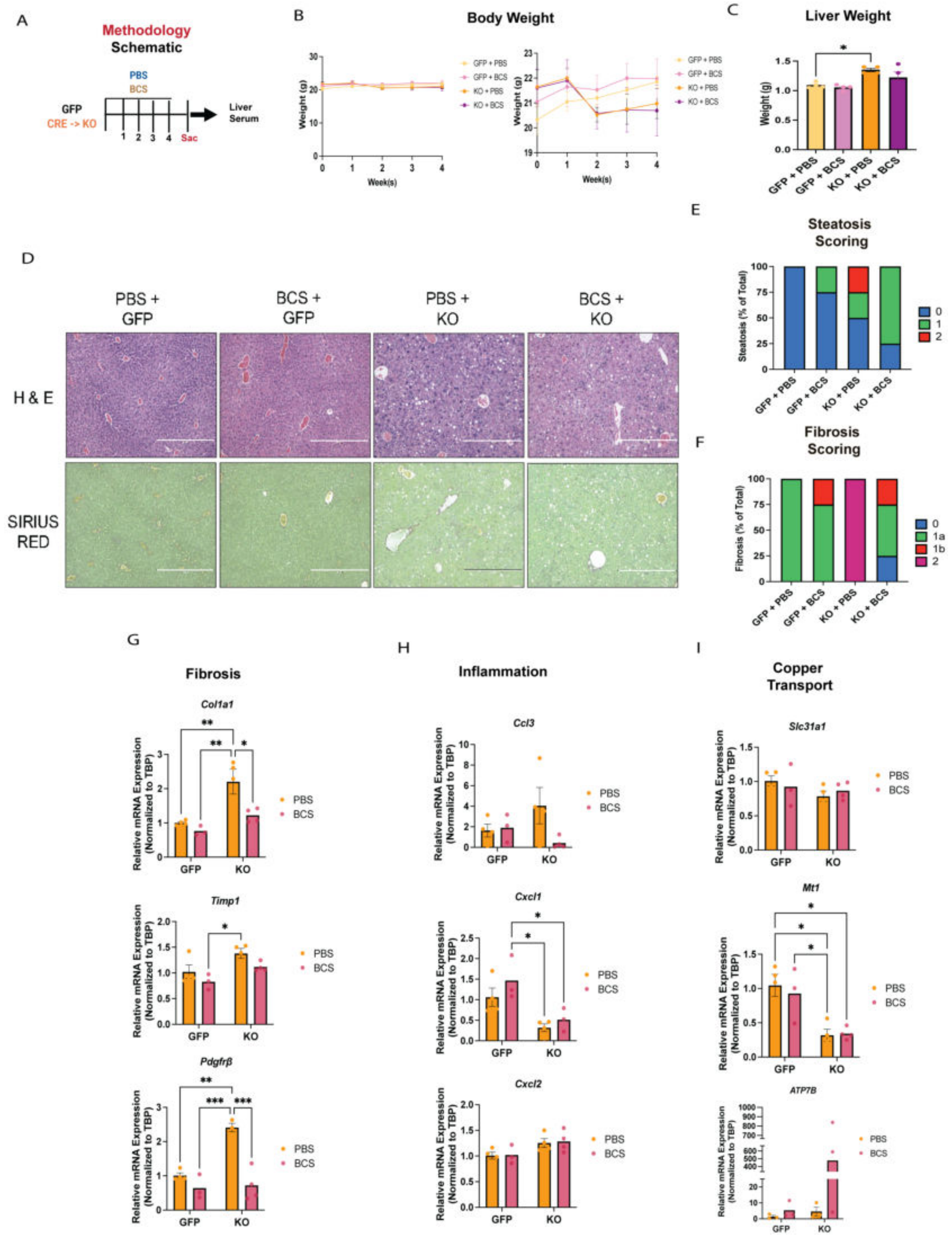


**(A)** – Schematic of Study Design and Model. **(B)** – Body weight across 4-wk BCS treatment period. **(C)** – Liver weight after 4-wk BCS treatment. **(D)** – Representative Histology. n = 4. **(E)** – Blind fibrosis, and steatosis scoring **(F)** in control vs. genetic mouse MASH livers post PBS or BCS administration for 4-wks. n = 4. **(G)** – **(H)** - RT-qPCR of pro-fibrogenic, and pro-inflammatory gene expression in control vs dietary mouse MASH livers +/- BCS. **(I)** – RT-qPCR of copper related gene expression in control vs genetic mouse MASH livers +/- BCS. n = 4. Data are shown as mean ± SEM. *P < 0.05; **P < 0.01; ***P < 0.001; NS, no significance, P > 0.05. Scale bars: 400 μm.

### Supplemental Figure 8

#### Copper Supplementation Does Not Restore Other Dysregulated Metal Levels in MASH

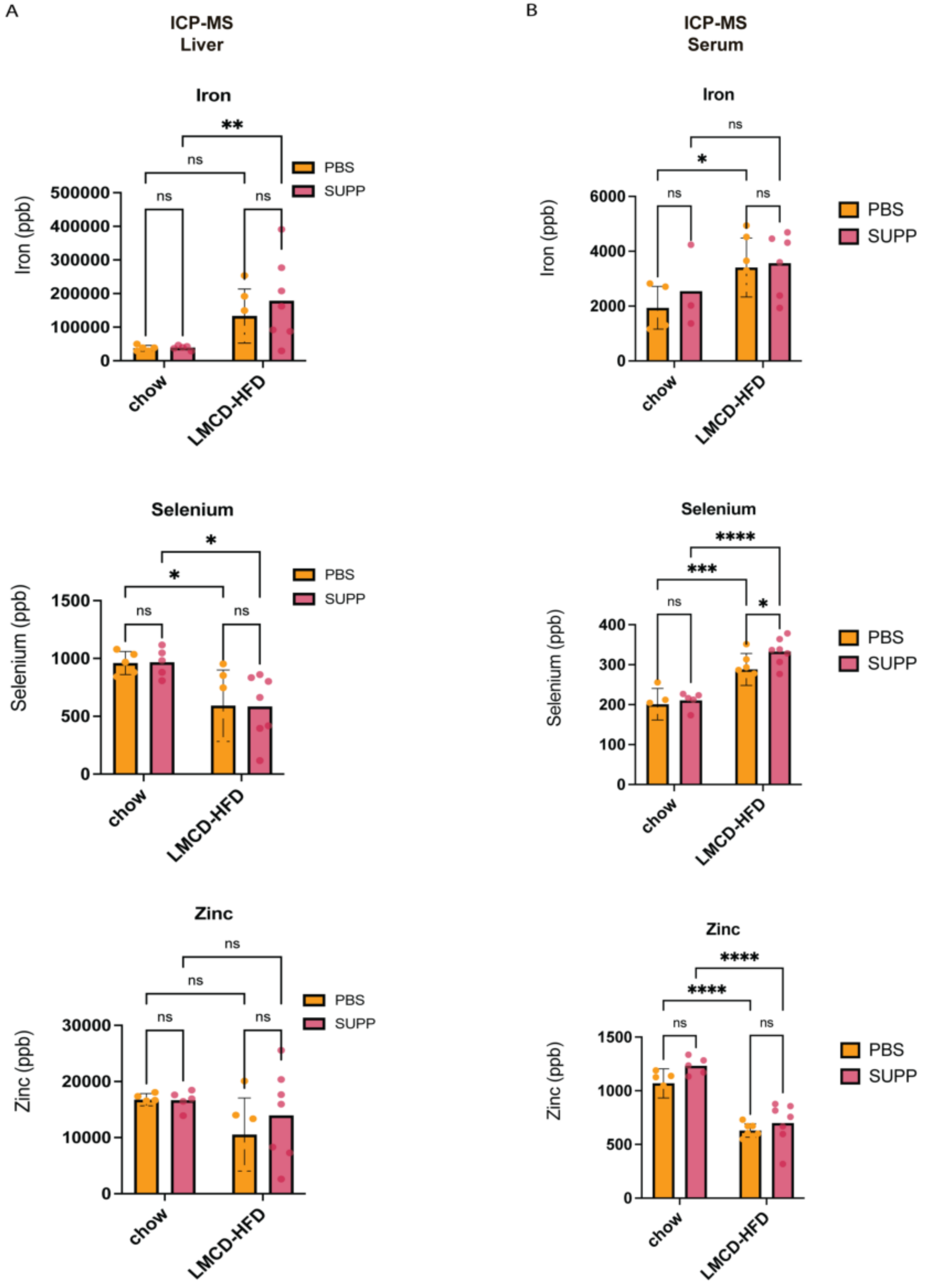


**(A)** ICP-MS showing normalized $^{63}$Cu in livers of control vs. dietary mouse MASH mice post PBS or CuS administration for 4-wks with chow or LMCD-HFD in prevention model. n = 5-7. **(B)** ICP-MS showing normalized $^{63}$Cu in serum of control vs. dietary mouse MASH mice post PBS or CuS administration for 4-wks with chow or LMCD-HFD in prevention model. n = 5-7.

| Table 1 - Homogenization Buffer | | | |
|---|---|---|---|
| **Regent Name** | **Stock Concentration** | **Working Concentration** | **Notes** |
| $MgAc_2$ | 1 M | 225 µl | |
| $CaCl_2$ | 1 M | 750 µl | |
| Tris-HCl | 1 M | 750 µl | pH 7.4 or 8.0 |
| BSA | - | 0.75 g | 1% of total volume |
| Protease Inhibitor | 1 tablet | Dissolved in 2 mL | |
| RNA inhibitor | 40 Units per µl | 2 Units per mL | 150 Units in 75 mL |
| Glycerol | - | 750 µl | 10% of Total Volume |
| MilliQ Water | - | Up to 75 mL | |
| Total volume = 75 mL<br>**Detergent Free** | | | |

| **Table 2 - Lysis Buffer** | | | |
|---|---|---|---|
| **Reagent Name** | **Stock Concentration** | **Working Concentration** | **Notes** |
| $MgAc_2$ | 1 M | 225 µl | |
| $CaCl_2$ | 1 M | 750 µl | |
| Tris-HCL | 1 M | 750 µl | |
| BSA | - | 0.75 g | 1% of total Volume |
| Protease Inhibitor | 1 tablet | Dissolved in 2 mL | |
| Triton X-100 | - | 750 µl | Either |
| NP-40 | - | 750 µl | Either |
| MilliQ Water | - | Up to 75 mL | |
| Total volume = 75 mL | | | |
